# An intragenic FAT1 regulatory element deleted in muscular dystrophy patients drives muscle and mesenchyme expression during development

**DOI:** 10.1101/2022.09.14.507898

**Authors:** Nathalie Caruso, Angela K Zimmermann, Tarana Nigam, Celine Becker, Karelia Lipson, Françoise Helmbacher

## Abstract

Fat1 is an atypical cadherin playing multiple roles that influence tissue morphogenesis. During mouse development Fat1 is required to modulate muscle morphogenesis through complementary activities in myogenic cells, muscle-associated connective tissue, and motor neurons, ablation of which leads to regionalized muscle phenotypes. We previously identified copy number variants (CNV) deleting an intragenic conserved non-coding element (CNE) in the human *FAT1* locus, that were enriched among muscular dystrophy patients with symptoms resembling those of Facioscapulohumeral Dystrophy (FSHD), compared to healthy individuals. Since such deletions of a putative cis-regulatory element had the potential to cause tissue-specific depletion of FAT1, they were postulated to act as symptom modifiers. However, activity of this CNE has not been functionally explored so far. To investigate the possible regulatory activity of this *Fat1-CNE*, we engineered transgenic mice in which it drives expression of a bi-cistronic reporter comprising the CRE-recombinase (Cre) and a myristilated-tdTomato fluorescent protein. The tissue-specific pattern of cre and tomato expression indicates that this enhancer has bipotential character, and drives expression in skeletal muscle and in muscle-associated mesenchymal cells. We extended our analysis of one of the transgenic lines, which exhibits enhanced expression in mesenchymal cells at extremities of subsets of muscles matching the map of *Fat1*-dependent muscles. This transgenic line exhibits highly selective CRE-mediated excision in scattered cells within the Tomato-positive territory hotspots. This represents a novel tool to genetically explore the diversity of muscle-associated mesenchymal lineages.

## Introduction

Conserved sequences in the non-coding genome regulate the tissue-specific expression of developmental genes. Their functional implication in recruiting transcription factors to ensure adequate gene expression and function constitutes a selection pressure that has guaranteed their conservation across evolution [1, 2]. While enhancer sequence variations, by modifying transcription factor recruitment, represent drivers of phenotypic diversity across evolution [2, 3], genetic abnormalities that disrupt functions of enhancers can also have pathological consequences and be at the root of genetic pathologies [4–6]. Furthermore, genome changes in non-coding sequences can also affect gene regulation by interfering with 3D chromatin architecture, thus impacting the regulation of large sets of genes within topologically associated domains (TADs) [7, 8]. Even though identification of disease causing mutations initially only focused on coding sequences, the availability of genome mapping techniques allowing the identification of small deletions or duplications of non-coding sequences [9, 10] or the mapping of long-range 3D genome interactions and their alterations [11], has expanded the tools to identify genetic causes of inherited pathologies. Given the modular distribution of tissue-specific enhancers in charge of distinct expression domains for a same gene, enhancer deletions can suppress portions (in space and/or time) of genes expression patterns while preserving expression in other domains/at other stages, resulting in phenotypes that partially reproduce the null phenotype.

While studying functional implications of the *Fat1* Cadherin gene in muscle morphogenesis [12, 13], and exploring the possible links [13–15] with Facioscapulohumeral dystrophy (FSHD), a human muscular dystrophy affecting restricted groups of muscles, primarily in the face and shoulder, we previously identified cases of copy number variants (CNVs) deleting a putative *FAT1* enhancer in patients with FSHD-like symptoms [13]. Fat1 is an atypical cadherin, involved in regulating tissue growth, morphogenesis and polarity during development [16–18]. Disrupting *Fat1* functions in mice interferes with morphogenesis of several organs, including kidney [19–21], eye and lens [21–24], Neural tube and brain [25], or muscle [12, 13]. Owing to a varying degree of redundancy between *Fat1* and the other family members [20, 25, 26], and a sensitivity to subtle differences in genetic background, *Fat1* deficiency causes phenotypes of varying severities, ranging from cases of severe congenital malformations such as kidney agenesis or glomerulotubular nephropathy, cyclopia, exencephaly, anophtalmia or coloboma, to milder phenotypes impacting tissue functions or homeostasis in adult mice [13, 21, 22, 25]. Aside from muscle, *FAT1* functions have also been linked with other human diseases [27], including autism [28], Kidney or eye pathologies [19, 23], and Cancer [29, 30]. These various functions of *Fat1* (or likewise of other Fat family members) rely on complementary expression and activities in different cells types, such as neurons, glia, kidney stroma or collecting ducts, smooth muscle cells, skin epithelium, but also myogenic cells, connective tissue, or tendons [12,20, 22, 29, 31–33].

Thus, with tissue-specific functions associated with phenotypes in multiple organs, *Fat1* is a typical example of gene for which enhancer deletions may disrupt part of the expression pattern, inducing modular phenotypes. In this context, the CNV deleting a putative *FAT1* cis-regulatory element identified in patients with muscle symptoms matching a subset of *FAT1*-dependent territories [13] have the potential to alter *FAT1* expression in the corresponding parts of its expression domain, with relevance to the specific symptoms. To begin to functionally explore this possibility, we have investigated the capacity of this putative enhancer to drive tissue-specific expression in FSHD-relevant domains in vivo, by producing transgenic mice with dual reporter modalities (CRE and myr-tdTomato). The tissue-specific pattern of cre/tomato expression indicates that this enhancer has bi-potential character, and drives expression in both muscle and mesenchymal tissues. We extend our description of one of the lines exhibiting a mesenchymal bias, to illustrate how the highly selective CRE-mediated excision in scattered cells within *Fat1*-expressing territories represents a novel tool to genetically explore the diversity of muscle-associated connective tissues.

## Results

### FSHD-associated Copy number variants delete a putative *FAT1* enhancer

We previously reported that copy number variants (CNVs) deleting portions of a Conserved Non-coding Element (CNE) in the human *FAT1* gene, were enriched among patients with FSHD/FSHD-like symptoms, compared to healthy individuals [13, 15]. This finding came from a Comparative Genomic hybridization (CGH) screen, using DNA arrays encompassing the 4q35 area, including *FAT1*, that had been conducted in a series of control individuals or patients with muscular dystrophy symptoms, which had been characterized by genetic diagnosis as either classical FSHD1 or as FSHD-like patients as they were not carrying the classical pathogenic *D4Z4* contraction and *DUX4* activating context [13]. Such *FAT1* CNVs were later also found in a small group of FSHD2 patients in which lowered *FAT1* levels correlated with phenotype severity [15]. This CNE, which included a large part of *FAT1* intron 16, exon 17, and extended over a portion of intron 17, matched the position of chromatin mark peaks (enriched in H3K27me3 and H3K4me3) from the ENCODE database (Figure 1A, Figures S1, S2), making it a good candidate to exert regulatory activity and act as tissue-specific enhancer. According to the ENCODE database, this element was indeed predicted to behave as a strong muscle enhancer (see ref [13] and Figure S1). This led to postulate that such deletions of a putative cis-regulatory element could result in tissue-specific (potentially muscle-specific) depletion of *FAT1*, if both alleles were affected, and may act as symptom modifiers, if one such heterozygous CNV co-occurred with an FSHD context. However, a formal assessment of the cis-regulatory activity of the deleted sequence was so far still lacking.

**Figure 1:**
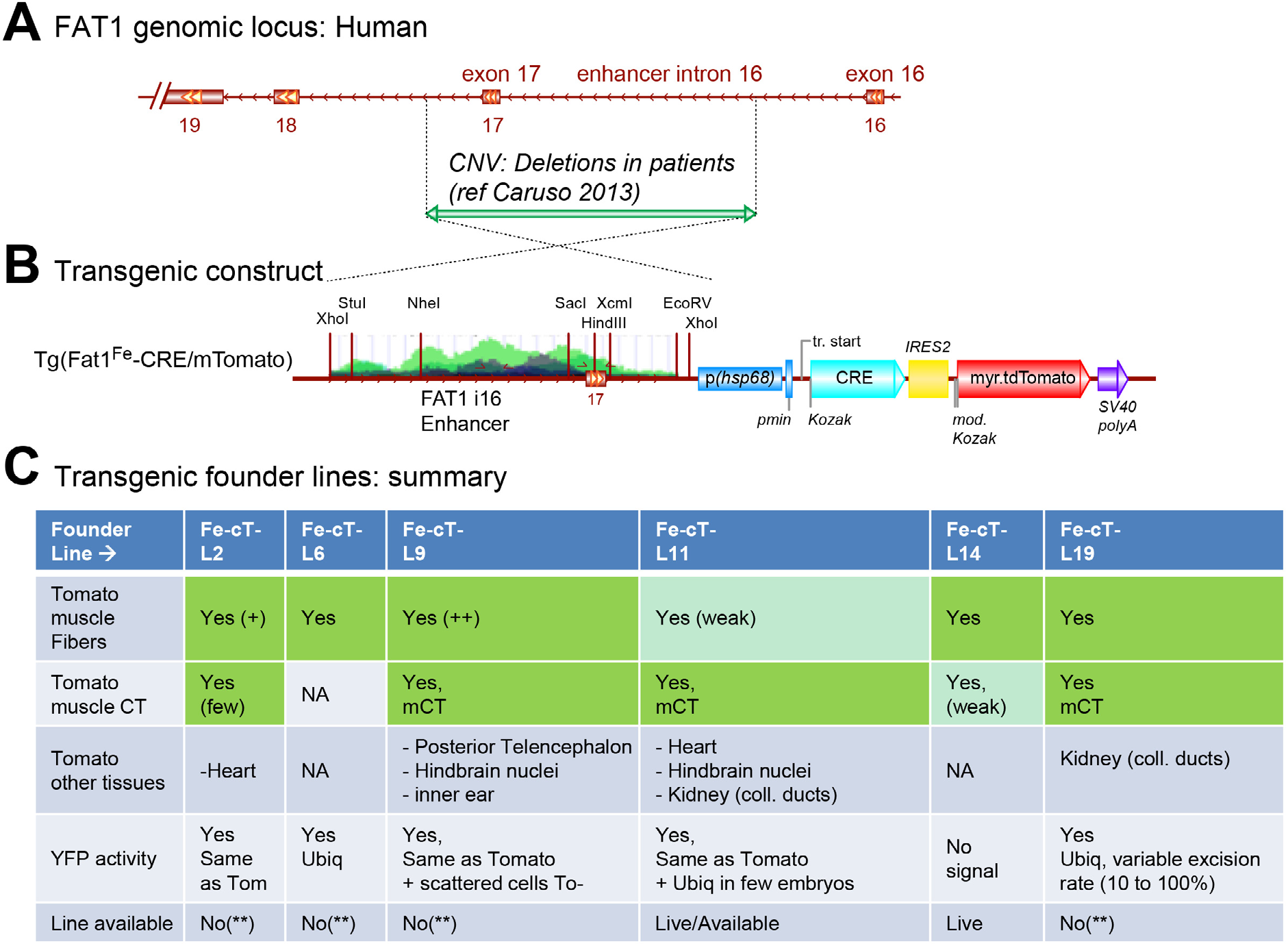
Position of previously identified copy number variants in the FAT1 genomic locus associated with FSHD. (A) Scheme of the human FAT1 locus showing exons 16 to 19, encompassing the enhancer cloned (*FAT1^Fe^* for “*FAT1*-FSHD-enhancer”), which matches the position (green arrowed bar) of deletions corresponding to copy number variants (CNVs) identified in FSHD patients (details in Ref [13] and in Figure. S1). (B) Design of the transgenic construct used for mouse transgenesis: the *FAT1^Fe^* CNE, followed by pHsp68 promoter, drives expression of CRE-IRES2-myr-tdTomato. (C) Summary of the expression patterns and cre-mediated activity profiles observed in the 6 founder lines analyzed *(Tg(FAT1^Fe^-cre/mTomato)Ln*, abbreviated as FF-cT-Ln, the n corresponding to the founder number). Transgenic embryos also carried the *R26^YFP^* reporter. Results were obtained upon immunostaining of cyro-sections, by following expression of Tomato with an anti-RFP antibody, and *R26^YFP^* expression with an anti-GFP antibody.

The genomic landscape around the *FAT1* gene contains multiple putative enhancers with predicted activity in muscle-relevant cell types (Figure S1). Focusing on the *FAT1-i16-i17* region deleted in the FSHD-associated CNVs (referred to as *FAT1^Fe^*), sequence comparisons and transcription factor binding sites (TFBS) searches highlighted the presence of several conserved TFBS including LEF/TCF, p53 and MEF3/Six1-4 (Figure S2). The FSHD-associated CNVs deleted either the full span or portions of the *FAT1-i16-i17* region [13]. These CNVs were present in a small percentage of healthy individuals (around 5%), indicating that heterozygous loss of the *FAT1^Fe^* CNE does not cause any pathology. However, their enrichment in FSHD1 and FSHD-like patients, in whom we also reported changes in *FAT1* expression levels [13, 15], suggested that the *FAT1^Fe^* CNE might indeed participate in the normal regulation of *FAT1* expression as postulated. Of note, this sequence also includes exon 17, implying that CNVs, by removing an exon, may also interfere with the protein sequence. However, exploring the possible consequences of exon 17 deletion will be the subject of future studies.

### Production of “FF-cT” mice

To determine whether the *FAT1^Fe^* CNE indeed exerts cis-regulatory activity in vivo, we cloned a sequence encompassing the full ENCODE peak. We engineered a transgene (called *Tg*(*FAT1^Fe^-cre/mTomato*), and abbreviated *FF-cT*) in which the *FAT1^Fe^* CNE is placed upstream of a robust promoter (hsp68 promoter, used for multiple in vivo studies, as it does not lead on its own to reporter expression without tissue-specific enhancers), to drive expression of a bi-cistronic cassette allowing dual expression of the CRE recombinase and of a fluorescent reporter myr-tdTomato (mTomato), a membrane-targeted red fluorescent protein (Figure 1B). The bi-cistronic character is conferred by the insertion of an internal ribosome entry site (IRES) between the two ORFs. The transgene design allows determining the tissue-type in which this putative enhancer is active by following mTomato expression, and simultaneously performing permanent CRE-mediated labeling of cells in which the transgene has been active, using CRE reporters such as *R26^YFP^* [34]. This construct was injected in mouse oocytes to produce transgenic founders; 7 adult founders (1 male, 6 females) were obtained. All founders were bred to wild type mice. Whereas one founder never transmitted the transgene to its progeny, the 6 other lines were successfully derived. Their properties were characterized by producing for each line, embryos carrying the transgene and the *R26^YFP^* reporter [34], and analyzing Tomato and YFP expression (results summarized in Figure 1C).

### Characterization of FF-cT transgenic lines

To uncover the tissue-specificity encoded by the *FAT1^Fe^* CNE, a first step involved screening reporter expression in all founder lines. We first analyzed mTomato expression in embryos collected from each of the 6 transgenic lines (identified as *Tg*(*Fat1^Fe^-cre/mTomato*)*Ln*, abbreviated as *FF-cT-Ln*, with n representing the line number), focusing on E12.5 embryos analyzed by immunohistochemistry on transverse sections. A summary of this analysis is provided in Figure 1C. We also compared mTomato expression with that of a *Fat1^LacZ^* allele, which reproduces endogenous *Fat1* expression pattern (previously described in refs [12, 13, 15]). This comparison is shown on an embryo carrying simultaneously the *Fat1^LacZ^* allele, expression of which was visualized by staining with Salmon Gal or anti-β-galactosidase antibodies, and the *FF-cT-L19* line (Figure 2), for which mTomato expression was followed with anti-RFP antibodies.

**Figure 2:**
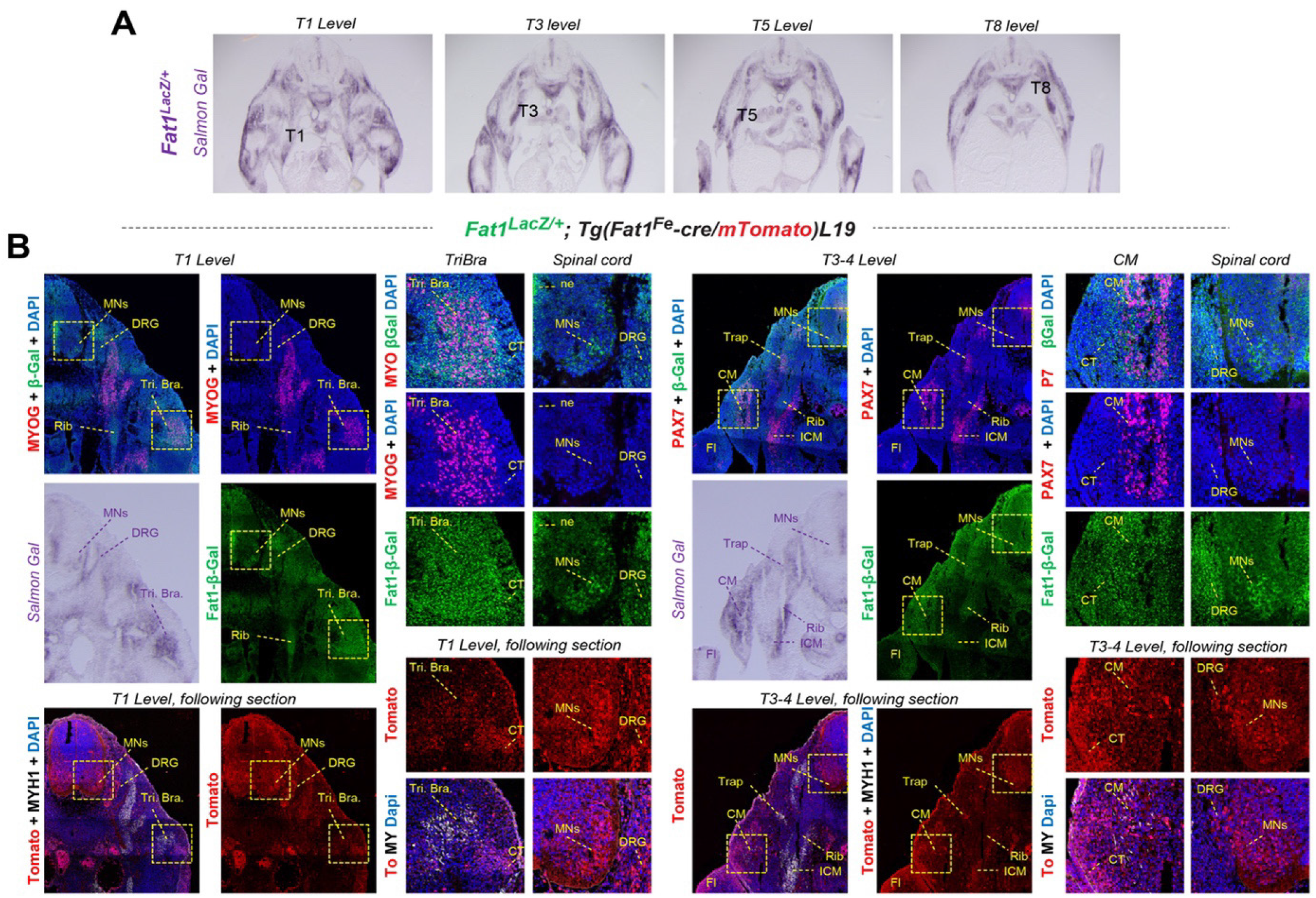
Comparative expression of *Fat1^LacZ^* and one of the *FF-cT* enhancer lines. (A) Salmon Gal staining of 4 cyrosections of an E12.5 *Fat1^LacZ^* embryo at different trunk levels, identified by the rib number. (B) immunohistochemistry analysis on cryosections at two levels of a *Fat1^LacZ^; Tg(Fat1^Fe^-cre-mTomato)L19* embryo at E12.5. The upper panels are stained with antibodies against beta-galactosidase (green), Myogenin (red, left/T1 level), Pax7 (Red, right/T3 level), and DAPI. The salmon Gal staining was done on the immediate neighbouring section. Bottom panels are stained with antibodies against RFP (Tomato, red), anti-MyhI (MHC, white), and DAPI. Square panels show higher magnification images of the regions indicated with dotted lines, focusing on a muscle (Triceps Brachii, T1 level, CM muscle, T3 level), or on the ventral spinal cord.

Overall, the *FAT1^FE^* CNE consistently drove expression in tissues that are part of *Fat1* expression domain. As summarized in Figure 1C, mTomato expression was detected at varying intensities in skeletal muscle in all the lines (6/6 lines), and in muscle-associated connective tissue (mCT) in the limbs, trunk or near shoulder muscles in several of the lines (5/6 lines). The *FF-cT-L19* line, shown for comparison with *Fat1^LacZ^*, drives mTomato expression in a combination of myogenic cells and mCT, and also reproduces expression in spinal motor neurons (Figure 2). Other lines only express Tomato in subsets of these components: some lines exhibited stronger expression in muscles (L9 (Figure 3), L2 and L6 (Figure S4), or L14, (not shown)); whereas one line (L11) exhibited a strong bias towards mCT expression (although myogenic expression is present, but at much lower intensity), with intense expression in clusters of mCT at the extremity of selected subsets of muscles (Figure 4, Figure 5) and low expression in myh1-positive myofibers. The high specificity of this expression pattern in mCT subsets, also part of *Fat1^LacZ^* expression domain (Figure 5), led us to explore this line (L11) in more detail (see following paragraphs). Finally, besides expression in the skeletal muscle system, these lines also exhibited Tomato expression in other tissues such as kidney, heart, or brainstem nuclei (Figure 1C, Figure S5), with at least 2 lines exhibiting each of these patterns.

**Figure 3:**
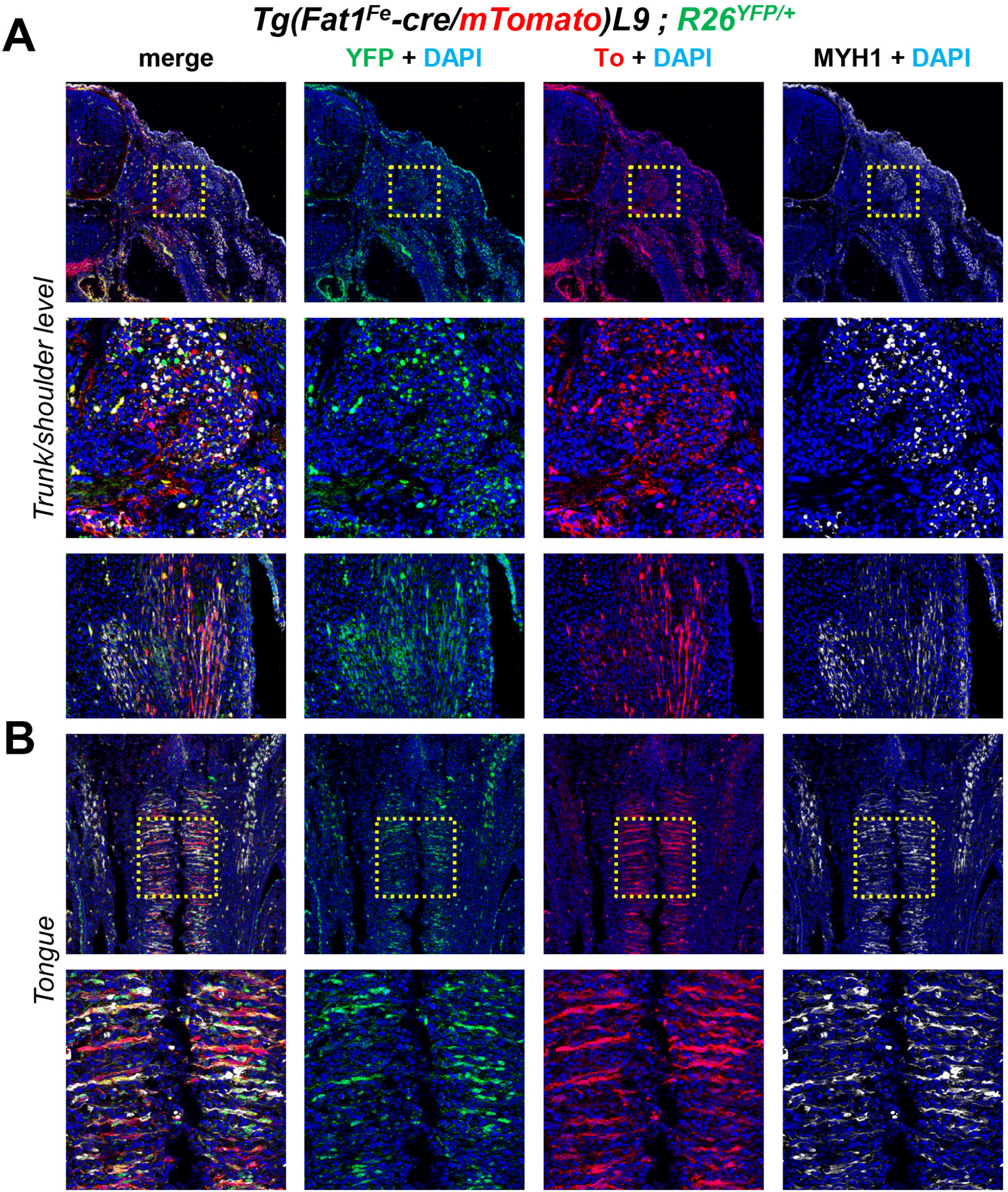
Example of founder line (FF-cT-L9) with robust expression and CRE activity in skeletal muscle fibers. Immunohistochemistry was performed on sections of E12.5 embryos carrying the line *Tg(Fat1^Fe^-cre-mTomato)L9*, combined with R26-YFP, with anti-Tomato (RFP, red), anti-MyhI (MHC, white), and anti-GFP (R26-YFP, green) antibodies.

**Figure 4:**
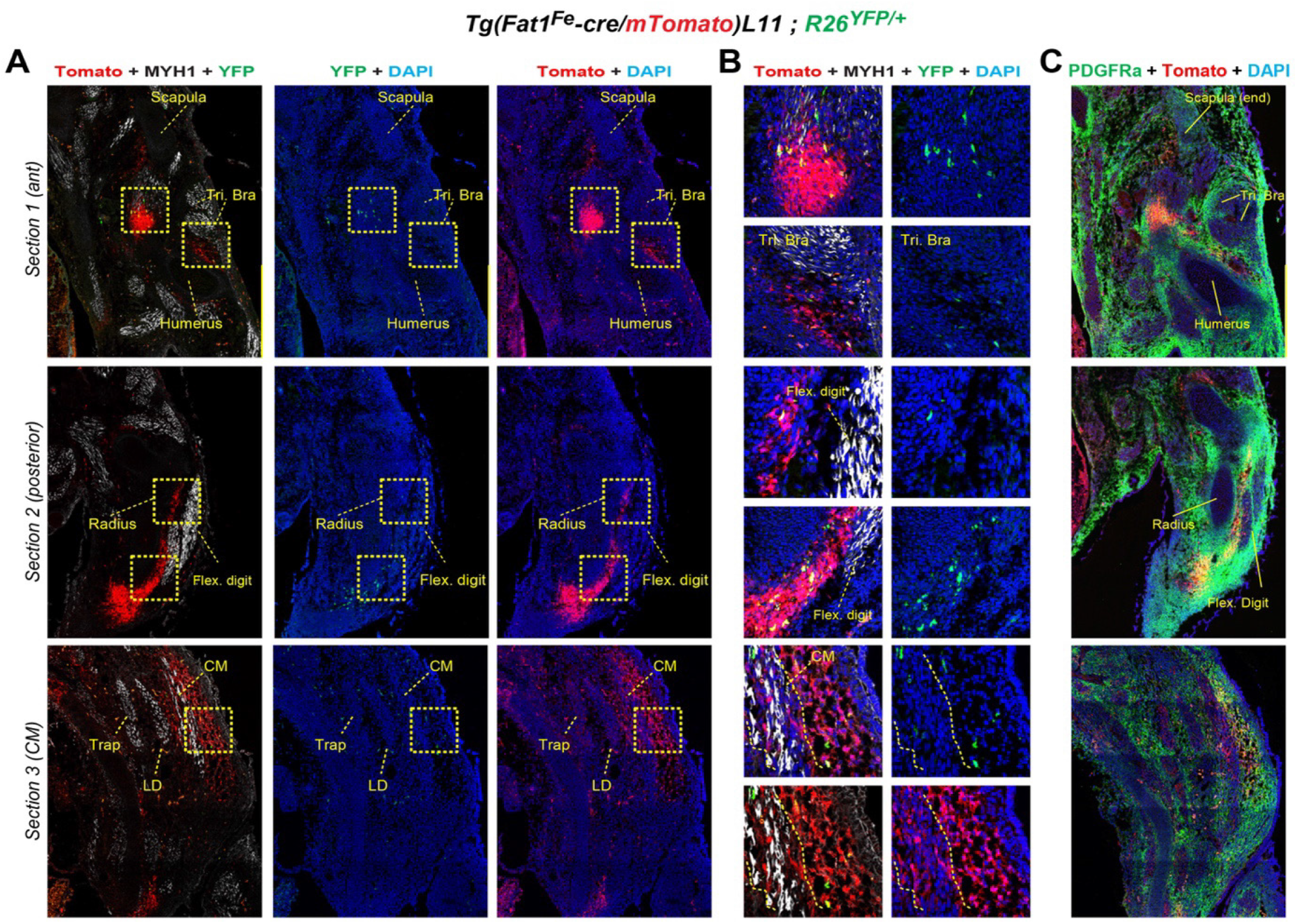
Example of a line *(FF-cT-L11)* with restricted expression and activity in selected subsets of mesenchymal cells at muscle extremities. Immunohistochemistry analysis of E12.5 embryos carrying the line *Tg(Fat1^Fe^-cre-mTomato)L11*, and *R26^YFP^*, with anti-Tomato (RFP, red), anti-MyhI (MHC, white, A and B), anti-GFP (*R26^YFP^*, green, A, and B), and Pdgfra (Green, C) antibodies, and with DAPI (blue). In contrast to other founder lines, Panels in (B) are higher magnification images of areas outlined with yellow dotted squares in (A). For each antibody combination, we imaged sections at three successive levels (section 1/row 1 corresponds to the shoulder level; section 2/row 2 to the forelimb level; and section 3/row3 to a thoracic level (half way through the CM muscle). In *FF-cT-L11* embryos, Tomato expression is low in muscle fibers, but high in specific hotspots of of muscle-associated mesenchymal cells at the extremity of some but not all muscles, whereas CRE-mediated activation of the *R26^YFP^* reporter occurs in few cells within the Tomato^+^ domain. All hotspots of high Tomato expression co-express Pdgfra.

**Figure 5:**
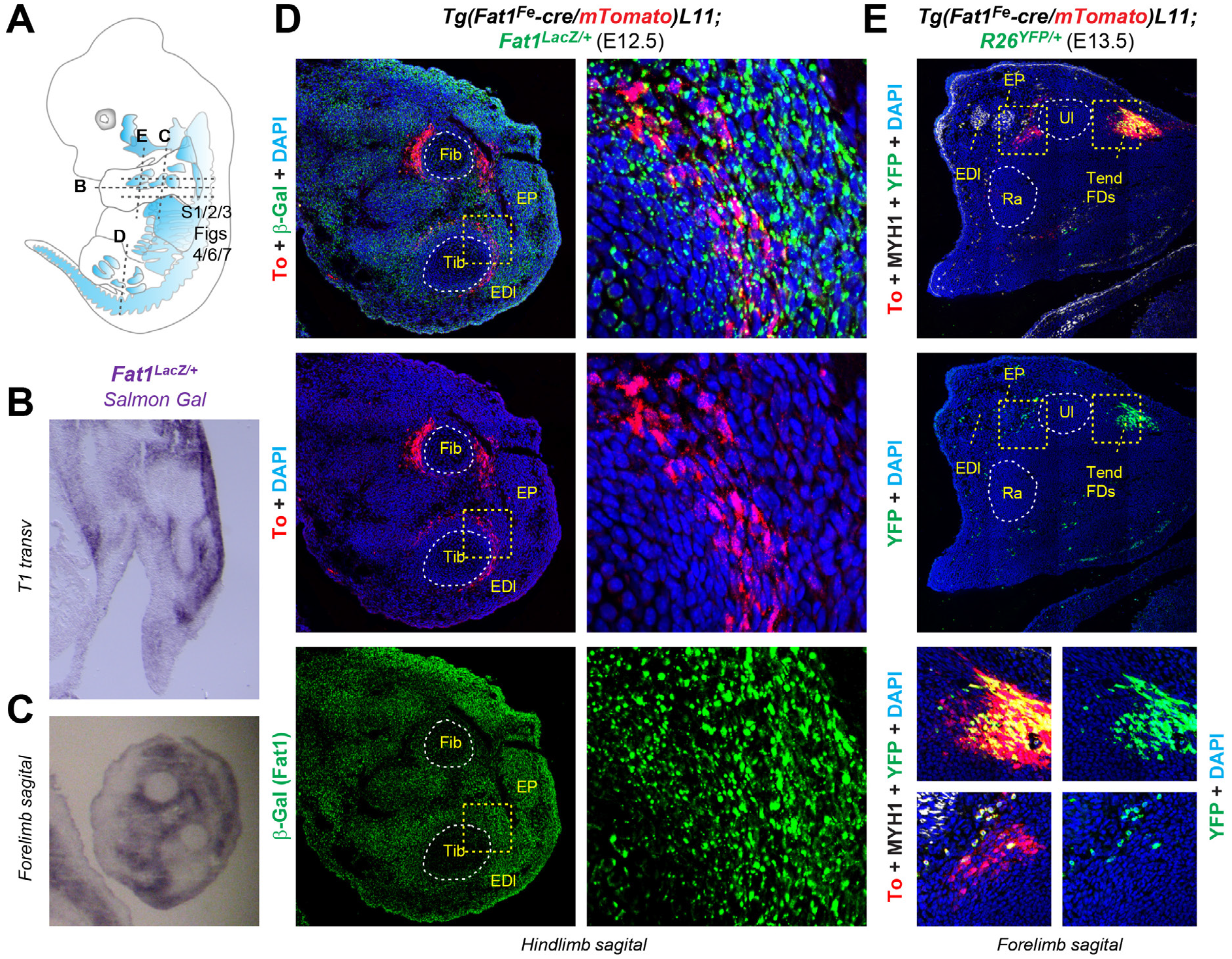
Mesenchymal hotspots of *FF-cT-L11* expression and activity at the bone muscle interface match sites of *Fat1^LacZ^* expression. (A) Scheme of an E12.5 embryo highlighting muscle organization, indicating the position and angles of sections shown in different following panels as well as the three successive levels shown in Fig. 4, 6 and 7. (B, C) SalmonGal staining of sections of *Fat1^LacZ/+^* embryos (orientations indicated in A). (D, E) Immunohistochemistry analyses of *FF-cT-L11* expression in comparison with endogenous *Fat1* pattern in an E12.5 *Fat1^LacZ/+:^ Tg(FAT1^Fe^-cre/mtdTomato)L11* embryo ((D), at hindlimb level), and with CRE activity in an E13.5 *Tg(Fat1^Fe^-cre-mTomato)L11; R26^YFP/+^* embryo ((E), forelimb level).

### Assessment of CRE-mediated recombination uncovers enhancer activity at early zygotic stages and restricted tissue-specific activity at mid gestation stages

The *Fat1^Fe^-cre/mTomato* transgene is designed to simultaneously visualize the tissue-type in which this enhancer is active (Tomato) and to induce permanent CRE-mediated recombination in these tissues. Sites of active CRE-mediated recombination can be visualized with the *R26^YFP^* reporter line [34]. We next assessed for each line the tissue-specificity of CRE-mediated recombination by crossing transgenic males with *R26^YFP^* carrying females (thus producing *FF-cT-Ln; R26^YFP/+^* embryos). Whereas some of the lines, exhibited a tissue-restricted pattern of YFP expression in a domain matching the observed profile of Tomato expression (Figure 3, Figure 4), others (in a varying percentage of embryos) exhibited ubiquitous YFP expression (Figure S3). While ubiquitous deletion was visible through direct YFP fluorescence at the moment of embryo collection (Figure S3B), the specific YFP patterns were analyzed by immunohistochemistry on sections, comparing them to Tomato expression (Figures 3, 4 and S4). The two patterns (Ubiquitous or specific) could be observed for a same line (even in a same litter), although the frequency of the ubiquitous pattern varied between lines. The ubiquitous pattern was observed for 4/6 lines (Figure S3). For CRE to have induced a ubiquitous *YFP* pattern, excision of the stop cassette must have occurred at very early stages (one to two cell stage depending on the percentage of recombined cells), indicating that the transgene (CRE and mTomato) is expressed at the 1-2 cells stage, supporting the possibility that *Fat1* may be expressed at these stages, consistent with its known expression in embryonic stem cells [35]. The co-occurrence of two YFP patterns in these lines was maintained after several generations, and we did not observe segregation over time of two different patterns of YFP expression, nor a progressive restriction of the pattern from generation to generation. This argues that the two patterns did not result from independent insertions in a same founder or the transgene in distinct chromosomal locations. Rather, the fact that a same line can exhibit two patterns in littermates likely reflected the stochasticity of the onset and extent of recombination, implying 1) that CRE expression levels at these early stages was low, only stochastically inducing recombination of the target locus in part of the Cre/Tomato-expressing cells, and 2) that this early wave of expression was transient and did not persist beyond the 1-2 cell stage. In embryos exhibiting a tissue-specific pattern, YFP expression was mostly included in the pattern of active Tomato expression at E12.5, with only a few scattered YFP+/Tomato-negative cells (indicating past transgene expression).

### Further characterization of the *FF-cT-L11* line

Although across all founder lines analyzed, expression in myofibers is well represented, supporting the idea that the enhancer is active in the myogenic lineage, we were particularly interested in the pattern exhibited by the *FF-cT-L11* line, in which expression/activity was highly enhanced in specific clusters of mesenchymal cells in the vicinity of some muscles, whereas Tomato expression was low (but not absent) in myogenic cells (Figures 4, 5, 6, 7). The spots of highest Tomato expression were found in the forelimb, at the distal extremity of the digit extensor muscles (Figure 4A, B, level 2), and in the shoulder area, near the upper side of the humerus, close to the upper insertion of the triceps brachii (Figure 4A, B, level 1). Tomato^+^ cells were also detected at the mesenchymal interface between the muscles and the nearest bone (triceps/humerus and digit-extensor/radius-cubitus interfaces in the forelimb, digit extensors/tibia-Fibula interfaces in the hindlimb (Figure 5)). Further caudally, a wider Tomato^+^area was detected around the cutaneous maximus (CM) muscle (Figure 4A, B, level 3), a subcutaneous muscle which emerges from the brachial plexus and extends subcutaneously towards the posterior flanks of the embryo, embedded in the skin. This muscle belongs to a particular group of skin-embedded muscles, recognized as panniculus carnosus, for which the muscle-skin interface is an alternative to the classical muscle-skeleton interface (whereas the other muscle extremity is classically inserted on bones). We previously showed that this muscle is particularly sensitive to *Fat1* loss of function, and that its expansion towards the skin requires mesenchymal *Fat1* activity, *Fat1* expression being particularly intense in the mesenchyme layer at the CM-skin interface [12, 13]. Similar to endogenous *Fat1* and *Fat1^LacZ^* expression, we observed a robust Tomato expression in *FF-cT-L11* transgenics at this CM-skin interface mesenchyme layer (both between the CM and the skin, and in more internal mesenchymal layers, between the CM, the latissimus dorsi (LD) and the trapezius (Trap.) muscles).

**Figure 6:**
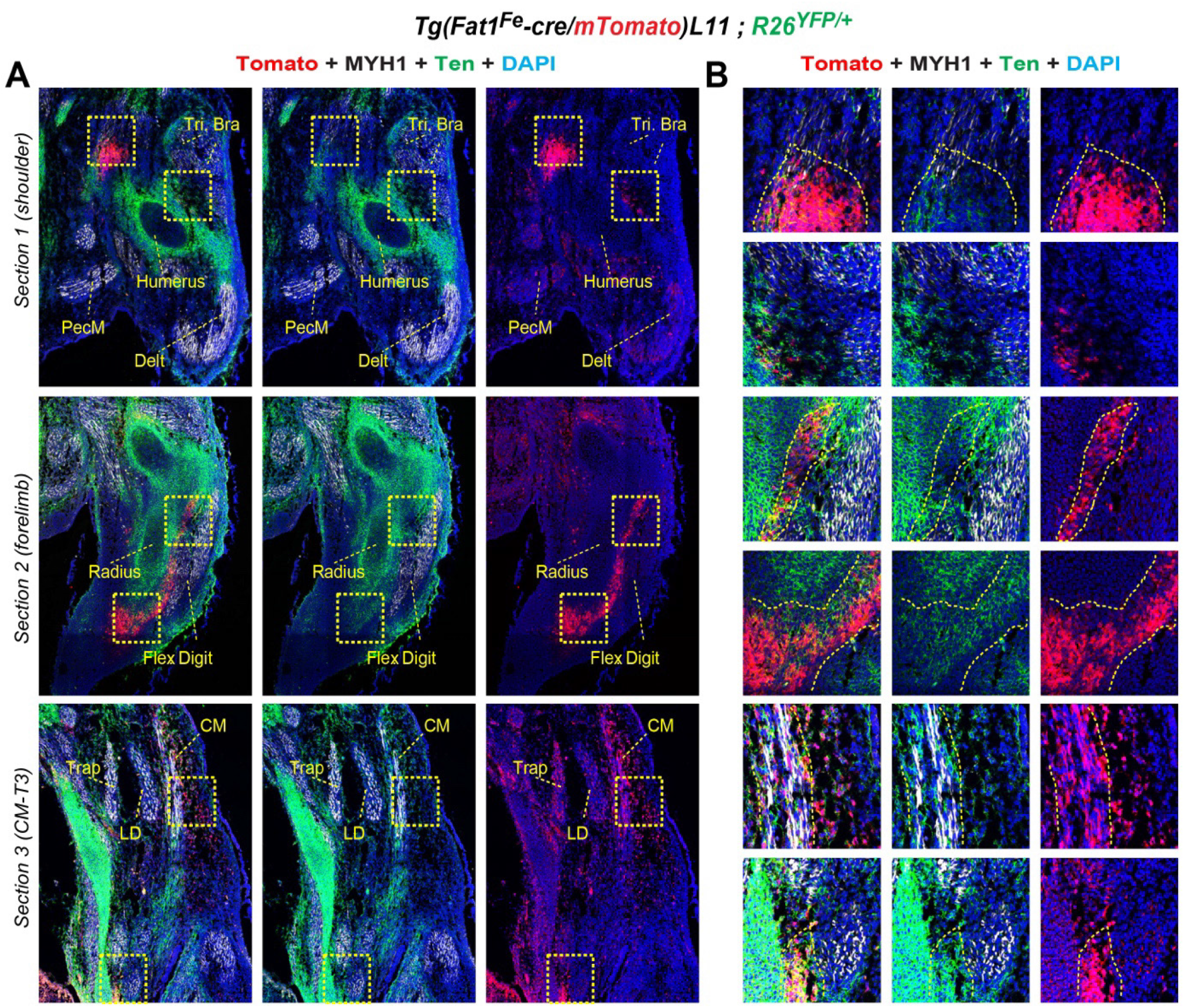
Comparison of *FF-cT-L11* expression with Tenascin C. Immunohistochemistry analysis of E12.5 embryos carrying the line *Tg(Fat1^Fe^-cre-mTomato)L11*, with anti-Tomato (RFP, red), anti-MyhI (MHC, white), anti-TenascinC (green), antibodies, and with DAPI (blue), at the same three levels as in Figure 4 (section 1/row 1 corresponds to the shoulder level; section 2/row 2 to the forelimb level; and section 3/row3 to a thoracic level (half way through the CM muscle). Panels in (B) are higher magnification images of areas outlined with yellow dotted squares in (A). All hotspots of high Tomato expression also coexpress TenascinC.

**Figure 7:**
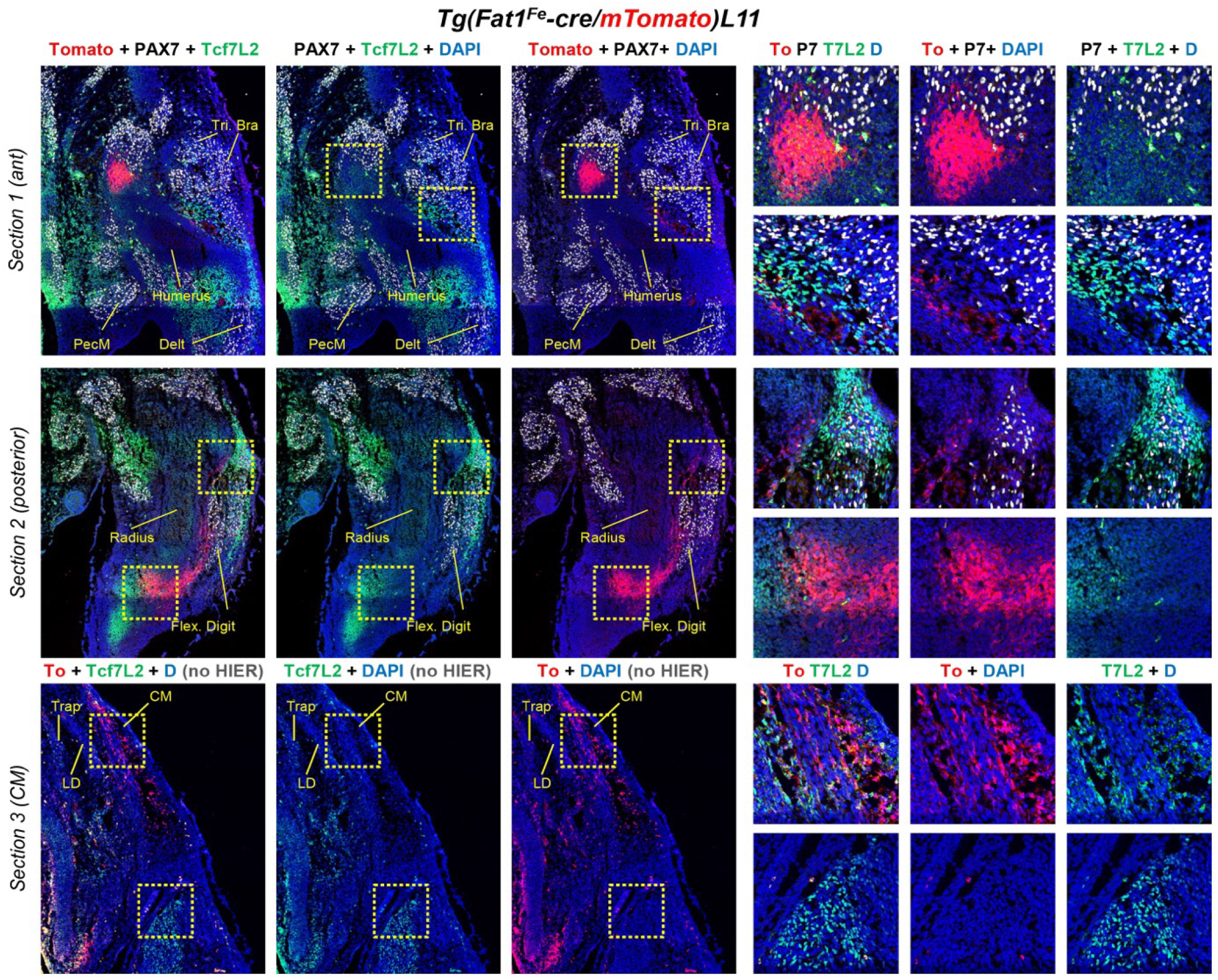
further characterization. (A) immunohistochemistry analysis of E12.5 embryos carrying the line *Tg(Fat1^Fe^-cre-mTomato)L11*, with anti-Tomato (RFP, red), anti-PAX7 (white), anti-TCF7L2 (green) antibodies, and with DAPI (blue), at the same three levels as in Figure 4 (section 1/row 1 corresponds to the shoulder level; section 2/row 2 to the forelimb level; and section 3/row3 to a thoracic level (half way through the CM muscle). Panels in (B) are higher magnification images of areas outlined with yellow dotted squares in (A). Hotspots of high Tomato signal do not co-express TCF7L2, whereas TCF7L2^+^ muscle attachment sites (or myogenic progenitor subsets) do not express Tomato.

In transgenic embryos (with a non-ubiquitous pattern of CRE-mediated *R26^YFP^* activity), YFP^+^cells were mostly concentrated in the Tomato^+^ hotspots, whereas only scattered/isolated YFP^+^ cells were seen in areas with lower expression levels or cell density. Whereas the proportion of YFP+ cells in the Tomato hotspots at muscle/skeleton interfaces was relatively important, only scattered cells were observed in the mesenchymal layer surrounding the CM muscle, and all of them were isolated from other YFP^+^ cells. The small number of YFP^+^ cells in this layer suggested that CRE-mediated excision had occurred/started at the stage of analysis, with limited clonal expansion (E12.5). None of the YFP^+^ cells lacked Tomato expression, arguing that (aside from the ubiquitous pattern reflected early activity/expression), cre-mediated deletion only occurred in cells with persistent Tomato^+^ identity. Although a low level of Tomato expression was detected in myofibers in some muscles (Figure 6, Figure 7), this was never associated with YFP expression at E12.5, arguing that transgenic expression is too low in the myogenic lineage in this line to achieve successful cre-mediated deletion. Nevertheless, it is interesting to note that this low level occurred in a specific set of muscles, matching our previous description of variable *Fat1* expression levels in muscles [13, 15]. Thus, CRE activity exhibits the same specificity and is entirely included in the domain of high Tomato expression. However, CRE activity is only sufficient to recombine the target locus in a very small proportion of the cells expressing the transgene. Altogether, this line represents an interesting tool to label subdomains of the *Fat1* expression pattern in mesenchymal cells surrounding muscle subsets affected by *Fat1* deletion, and to carry-out clonal analyses of recombination events by following YFP-positive cells.

In the areas with high *FF-cT-L11* transgene expression, Tomato^+^ cells expressed the mesenchymal marker Pdgfra (Figure 4C), and TenascinC (Figure 6), an extracellular protein produced by connective tissues including bones and tenocytes. Although this profile is typical of tendon attachment sites, it did not highlight every muscle extermity: Double labelling with Tomato and Tcf7L2 (previously called Tcf4), a transcription factor known for its expression in tissue fibroblasts marking muscle extremities [36, 37], uncovered a complementarity between the two patterns, with overlap in exceptional positions (Figure 7). An emblematic example is found in the limbs, where a same muscle (digit extensor) has a proximal extremity marked by Tcf7l2, and a distal extremity marked by *FF-cT-L11* expression. Likewise, the connective tissue surrounding the CM muscle (towards which the CM expands) expressed Pdgfra and TenascinC, but not Tcf7L12 (Figure 4C, Figure 6, Figure 7). These findings illustrate the emerging notion of a molecular heterogeneity of muscle-associated mesenchymal cells, and shows that such cells at muscle attachment sites can even differ between the two extremities of a same muscle.

## Discussion

In the present study, we confirm that the sequence deleted in FSHD-associated CNVs is capable of exerting transcriptional regulation activity in vivo, by driving reporter expression in developing muscles and in muscle-associated connective tissue cells at muscle attachment sites. These expression domains match the parts of *Fat1* expression domain that were shown to be required for the modulation of muscle morphogenesis [12, 13, 15].

### Muscle-relevant activities of the *FAT1* enhancer in myogenic and mesenchymal cells

Our previous work has illustrated how the control of muscle morphogenesis by *Fat1* involves distinct functions in several muscle-relevant cell types, including myogenic cells, but also motor neurons, and mesenchymal cells, [12, 13]. Muscle growth occurs via expansion of a progenitor pool, the commitment of cells towards myogenic differentiation, the subsequent progression along a well-characterized differentiation trajectory, and the fusion of resulting myocytes, to form multinucleated myofibers in charge of the contractile function [38, 39]. Subsets of muscles also involve a step of muscle progenitor migration, with limb muscles originating in somites, while head and neck muscles derive from cardio-pharyngeal progenitors [40, 41]. Although *Fat1* inactivation in myogenic progenitors (with *Pax3-cre*) does not prevent muscle growth per se, it interferes with the polarized myoblast migration, leading to the dispersion of myoblasts in ectopic positions in the limb [13]. Whereas the control of myogenic growth and muscle migration rely on events intrinsic to the myogenic lineage, muscle morphogenesis is also highly dependent on signals from non-myogenic cells derived from the mesenchymal lineage (reviewed in [38, 42]). Along this line, whereas muscle-specific *Fat1* ablation does not reproduce the full span of muscle phenotypes observed in constitutive knockouts [13], ablation in the lateral-plate-derived mesenchymal lineage, driven by Prx1-cre or the inducible Pdgfra-CRE/ERT line, was sufficient to reproduce the specific pattern of shape alterations (aberrant muscle attachment sites, and mis-oriented myofibers) in subsets of limb muscles and the failure of CM muscle expansion, whereas facial muscle phenotypes were reproduced by ablation in the neural crest lineage (Wnt1-cre) [12]. The latter phenotypes include the appearance of ectopic muscles, of aberrant attachment sites, and the formation of muscle bundles with abnormal orientations. These phenotypes are reminiscent of phenotypes resulting from disrupting transcription factor genes acting in subsets of muscle-associated CT mesenchymal cells, such as Osr1 [43], Tbx3 [44], Tbx5 [44] [45], Tcf7l2 [36, 37]. Actions of these transcription factors involve the regulation of CT-derived ECM or ECM-interacting proteins that influence myogenesis [43, 45], even though for each of these genes, the muscle groups in which they act differ. This highlights the emerging notion of the existence of multiple subtypes of muscle-associated mesenchymal progenitor lineages, distinguished from each other by their molecular, regional and possibly functional characteristics [46]. This notion is supported by recent single cell RNA sequencing studies, which also hint towards the existence of similar molecular diversity in adult [47–50] but also embryonic [51] muscle-resident mesenchymal progenitor cell subtypes.

Our finding that a *FAT1* enhancer exhibits dual transcriptional tissue-specificities, encompassing myogenic and mesenchymal lineages, is in line with the complementary activity of *Fat1* in both cell types. Interestingly, although encoded by the same enhancer, the mesenchymal and myogenic components can be uncoupled from each other, with line 9 exhibiting a myogenic bias, whereas the line 11 exhibits a mesenchymal bias. Even if the reason for such uncoupling is not known (it may likely result from repressor action of flanking genomic sequences in the loci in which the transgene was inserted, which differ between founder lines), the lines in which these two components are separated illustrate the modularity of *Fat1* regulation, and allow dissecting each of them in more detail. It is particularly intriguing that the line with mesenchymal bias shows a pattern of expression highlighting regional subsets of muscle-associated mesenchymal cells, complementary to that of Tcf7l2, another marker of subsets of muscle attachment sites. This highly specific expression matches the regions displaying overt muscle attachment phenotypes in *Fat1* mutants, providing a potential framework to explain regional specificities in FSHD. Furthermore, this transgenic line (L11) may represent a useful tool to better characterize and distinguishing this novel specific mesenchymal progenitor lineage.

### Relevance of *FAT1* enhancer deletion as putative FSHD modifier

Aside from the potential implication of *FAT1* in FSHD, the main cause of FSHD is a well described complex genetic abnormality on Chromosome 4 (4q35), which combines the suppression of a mechanism of epigenetic silencing of a gene encoding the transcription factor DUX4 (encoded by *D4Z4* macro-satellite repeats), with the presence of a polymorphism enhancing stability its RNA, resulting in enhanced production of the DUX4 protein, toxic for muscles [52–54]. *DUX4* is normally only expressed at early zygotic stages to activate transcription of the zygotic genome [55, 56], after which stages it is subject to a robust epigenetic silencing [57]. The loss in FSHD patients of this epigenetic silencing results in the most frequent cases (FSHD1), from the shortening of the *D4Z4* repeat array, and from consequent changes in topological genomic structure, affecting regulation of neighbor genes in the area [54, 57, 58]. In rarer cases (FSHD2), it occurs as a result of mutations in genes involved in establishing a repressive context via chromatin organization or DNA methylation (*SMCHD1*, and in rare cases *DNMT3B*) [59, 60]. Although the events involved in *DUX4* activation have been clarified, the identified mechanisms do not fully explain the highly selective topography of muscle symptoms. Furthermore, the severity of FSHD symptoms varies between patients, ranging from childhood onset to individuals remaining asymptomatic until advanced age in spite of a pathogenic DUX4-compatible context, implying the existence of genetic modifiers of disease severity [61–63], among which the first known were *SMCHD1* and *DNMT3B* [64, 65].

In this context, our findings that the mouse *Fat1* gene, which human homologue *FAT1* was localized in the vicinity of the FSHD locus, was required for muscle development [12, 13, 38], and that *Fat1* mutations in mice caused muscle phenotypes with a topography matching that of FSHD symptoms, originally suggested that *FAT1* dysfunction might potentially contribute to FSHD [13]. This idea was first supported by the identification of the CNVs studied here that deleted the putative *FAT1* enhancer, and were enriched among FSHD patients [13]. It was further supported by the identification of pathogenic *FAT1* single nucleotide variants (SNVs), not only in rare patients with FSHD-like symptoms not carrying *DUX4*-activating contexts [14], but also in classical FSHD1 patients [66], Most of these pathogenic *FAT1* SNVs were absent from snp databases of healthy individuals, and their overall frequency among FSHD-like patients was significantly higher than in the healthy population. Using a minigene splicing assay [67], 4 of these SNVs were shown to alter *FAT1* RNA splicing, causing either complete exon skipping or truncation of the mutated exon [14]. As in FSHD-like patients, these *FAT1* SNVs were carried by only one allele, they were postulated to act as dominant negatives [14]. Further support came from the observation of reduced *FAT1* RNA levels in muscles with early-onset symptoms in FSHD1 and FSHD2 patients [15]. The possibility that *FAT1* repression would result from *DUX4* overexpression, initially suggested by data from myoblasts transfected with exogenous *DUX4* [13], was ultimately ruled out by RNA interference studies silencing endogenous *DUX4* in FSHD myoblasts [15]. Instead, comparisons of 3D genome conformation between FSHD and control cells uncovered changes in long-range association between the *FAT1* locus and a region near the *D4Z4* array that were likely to impact the regulation of *FAT1* expression [58]. Collectively, the findings above raised the possibility that *FAT1* dysfunction, which on its own leads to FSHD-like symptoms in mice, and is uniquely found in rare human FSHD-like cases, co-occurs with FSHD, either as a regulatory consequence of disrupted chromatin regulation in FSHD1 and FSHD2, or as an association of *FAT1* variants with FSHD. While heterozygous variants may not be sufficient to cause symptoms, they may act as FSHD symptom modifiers by in particular when combined with a genetic context that also perturbs gene regulation in the 4q35 area by perturbing TADs.

However, the clinical presentation of FSHD symptoms only matches a subset of the tissue-types in which *FAT1* is known to exert its functions, sparing key organs such as kidney or brain, implying that the FSHD context must spare *FAT1* expression in these tissues. While at present, whether regulatory changes induced by the classical FSHD context are tissue specific is not known (most of them have been studied in myogenic cells only), genetic alterations of *FAT1* regulation, such as the deletion of putative cis-regulatory elements, have the potential to cause tissue-specific changes of *FAT1* expression in cell types relevant to muscle biology, providing a potential frame for the selective FSHD topography. Here we provide confirmation that the CNE deleted in subsets of FSHD patients is indeed capable of driving gene expression in mice in portions of *Fat1* expression pattern that are relevant to its function in muscle development, including developing myofibers, and muscle associated mesenchymal cells at muscle-tendon and muscle-skin interfaces. Analysis of transgenic expression outside of the musculoskeletal system uncovered only minimal expression in tissues or cell types that would account for *Fat1* functions in tissues not involved in FSHD. Although two lines exhibited kidney expression, this was exclusively in collecting duct epithelium, and not in stromal cells, where *Fat1/Fat4* play their major roles [20].

One way of assessing experimentally in mice the functional implication of this enhancer might have been to knock it out in mice and evaluate whether homozygous deletion would be sufficient to deplete *Fat1* expression in relevant cell types and to reproduce the muscle phenotypes that we described in muscle-specific and/or mesenchyme-specific *Fat1* mutants [12, 13]. However, our analysis of the genomic locus had revealed the presence of other conserved segments with predicted muscle-like regulatory activity (Figure S1), and were confirmed to be the sites of DNase hypersensitivity in myoblast cultures, although no clear differences were detected between FSHD-derived myoblasts compared to myoblasts from healthy patients [68]. Under the light of work discussed above, suggesting that the component of *FAT1* expression domain relevant to muscle morphogenesis, and possibly to FSHD symptoms, might occur in fibroblast/mesenchymal lineage rather than (or in addition to) in myoblasts [12, 38], it will be interesting to study in future whether the changes in *FAT1* regulation occur in mesenchymal/fibroblast lineage rather than in myoblasts. Furthermore, *FAT1* is located in a conserved genomic region, between *FRG1* and *SORBS2*, now identified as a TAD [35, 58], in which long-range interactions allow shaping gene regulation. Thus, the *FAT1* locus represents a typical situation in which shadow enhancers with similar tissue-specificity may cumulate their activities to ensure the robustness of the expression pattern [69, 70]. Several attempts to delete single enhancers in the cis-regulatory context of developmental genes with robust expression, have been shown to only modestly impact gene expression and function, owing to the presence of multiple shadow enhancers with redundant activities [69, 70]. However, such single enhancer deletions may nevertheless induce phenotypes in a sensitized background, in which other elements of a same genetic or regulatory cascade are also compromised [69, 70]. The association of the CNV deleting the *FAT1* enhancer with the genomic context of FSHD1 or two, both of which interfering with gene regulation in the 4q35 locus has the potential to constitute such a synergistic situation leading to *FAT1* dysregulation (in addition to enhanced *DUX4* expression).

Instead, owing to the dual reporter and CRE-mediated deletion, the transgenic lines described here were designed to be used as tool to indirectly model in mice the effect of conditionally ablating *Fat1* in tissues driven by the *FAT1* enhancer, ultimately aiming to ask if this causes FSHD-like phenotypes. However, given that CRE activity was only detected in a small fraction of Tomato expressing cells, we anticipate the impact of tissue-specific cre-mediated Fat1 deletion to be limited, only allowing assessing cell autonomy versus non autonomy of phenotypes, or for lineage tracing studies.

## Methods

### Construction of the Fat1-CNE-CRE-IRES2-myr-tdTomato transgene

A 1.7kb human genomic region spanning a large part of intron 16, *FAT1* exon 17, and a small part of intron 17 (coordinates >Human Mar. 2006 chr4: 187,763,947-187,765,500 (-) or >Human Feb. 2009 chr4:187,526,753-187,528,506), (this DNA fragment is subsequently referred to as *FAT1^Fe^*), flanked by gateway sites AttB3 and AttB4 compatible with a multicassette gateway system [71], was produced by gene synthesis and cloned into pBluescript (Eurofins, plasmid p1-Q426). This *FAT1^Fe^* element was then transferred in a vector engineered to contain a pHsp68 promoter, driving expression of a bicistronic gene encoding the CRE recombinase, and m-tdTomato (myristoylated tandem dimeric Tomato), the two separated by a modified IRES2 sequence. The cloning strategy to assemble all necessary elements and produce the final transgene involved the following steps: 1) The L3-L4 Gateway cassette from p1 plasmid was combined with a modified pDONR221 (containing AttP3-ccdB-CmR-AttP4 as described in [71]) via a BP clonase reaction, to produce a plasmid called pNC13 (AttL3-*FAT1^Fe^*-AttL4). 2) A synthetic CRE sequence flanked by AttB1-AttB2 sites was cloned (by Eurofins) upstream of a IRES2-DsRed-express2 cassette (plasmid pIRES2-DsRed-express2 from Clontech/SakaRa, #632540), producing a plasmid called p4 containing AttB1-CRE-AttB2_IRES2-DsRed-express2. 3) We next used another plasmid produced in the lab following the multicassette system startegy, which contained an AttR3-ccdB-cmR-AttR4 cassette, followed by a promoter from the pHsp68 gene [72], and inserted the AttB1-CRE-AttB2_IRES2-DsRed-express2 cassette, to produce a plasmid called pNC12. 4) The *FAT1^Fe^* sequence from pNC13 was transferred upstream of the hsp68 promoter in pNC12 via a gateway LR cloning reaction, producing a plasmid called pNC14 (AttB3-*FAT1^Fe^*-AttB4-pHsp68-AttB1-CRE-AttB2-IRES2-DsRed-express2). 5) The sequence of DsRed-express2 was subsequently replaced by mtdTomato sequence in the final vector, since a first attempt to produce transgenic founders with pNC14 had failed. To do so, we replaced in pNC14 an AvrII-NotI fragment containing part of IRES2 and DsRed2, with an equivalent fragment from a pIRES2-mtdTomato plasmid (kind gift of N. Denans), thus producing our final transgenic construct (pNC17).

### Sequence analyses to characterize the *FAT1^Fe^* conserved sequence

The sequence of the FSHD-associated CNVs we defined previously [13]. We used the Vista Enhancer browser to define the conserved region and obtain orthologue sequences from other species (updated coordinates were obtained via the USCS browser). Analysis of transcription factor binding sites was done using an online suite for genome analyses from Genomatix (now Precigen Bioinformatix, Germany), including MatInspector matrices and matrix matches in the conserved sequences.

### Generation of transgenic mice

The final construct was digested with NdeI and Sap I to isolate the transgene, which was gel purified according to recommended procedures for transgenesis. After purification, the transgenes were injected into pronuclei of fertilized eggs from B6/CBA mice as described previously. Injected eggs were implanted by injection into the oviduct of pseudo-pregnant foster mothers. Among the mice born, 7 positive founders were identified by PCR genotyping of tail DNA with several primer couples spanning the construct (5’ and 3’ sides, as well as interface between key elements). Genotyping oligonucleotides were the following: o.FF3 (Fw): 5’-TGA GCT TTT CCA TTG GCC TCT GTT GC-3’ ; o.phsp68-5 (Rev): 5’-TAG GAA CTA GAG GCT CTG TCC CAG C −3’. Alternatively, we also assessed the presence of other parts of the transgene with the following primers: o.phsp68-3 (Fw): 5’-GCG ATG ATC CCG TCG TTT TAC-3’; o.cre_1 (Rev): 5’-GGT TCT GCG GGA AAC CAT TTC −3’. Each of the founder (F0) mice (1 male, 6 females) was bred to wild type B6D2F1/JRj mice (Janvier labs, resulting from a cross between C57Bl6/JRj and DBA/2JRj mice) to obtain F1 transgenic carriers, and 6 lines were successfully derived and characterized (referred to as *Tg*(*FAT1^Fe^-cre/mTomato*)*n*, and abbreviated as FF-cT-Ln, with n corresponding to the founder number. Although most lines were maintained alive for the present characterization, most of them had to be sacrificed during the Covid pandemic (or were lost as a consequence of drastic restrictions), and only the FF-cT-L11 line remains available for further studies.

### Mice

#### Ethics statement

All procedures involving mice were in accordance with the European Community Council Directive of 22 September 2010 on the protection of animals used for experimental purposes (2010/63/UE), with the French law and with institutional guidelines for animal Research from institutional Ethics Committees for animal experimentation (of Marseille and of CNRS Campus orleans, respectively registered as N°14 and N° 003 by the French national committee of ethical reflection on animal experimentation). Transgenesis experiments were carried out at the TAAM mouse facility (TAAM, CNRS UPS44, Orleans), and mice were later transferred to and maintained at the IBDM mouse facility, under an agreement (Number D13-055-21) delivered by the “Préfecture de la Région Provence-Alpes-Côte-d’Azur et des Bouches-du-Rhône.

The mouse lines used in this study are the following: The FF-cT transgenic lines were generated as described above and are identified as *Tg*(*FAT1^Fe^-cre/mTomato*)*nFhel*, with n representing the line number (as summarized in Figure 1); A *Fat1^LacZ^* genetrap allele (*Fat1*^*Gt(KST249)Byg*^ [13, 73]); the CRE-reporter line *R26^YFP^* (Gt(ROSA)26Sor^tm1(EYFP)^Cos line (Jackson laboratory mouse strain 006148, [34]), a Wnt1^cre^; driver line (H2az2^Tg(Wnt1-cre)11Rth^, [74]) for comparison.

### Immunohistochemistry

Embryos/tissues were collected in cold PBS, fixed in 4% PFA (in PBS) for 3-4 hours on ice, rinsed 3 times in cold PBS, and cryoprotected by overnight immersion in a 25% sucrose solution in PBS. The samples were then embedded in PBS, 7.5% gelatin, 15% sucrose, and frozen by immersion in isopentane refrigerated in a carboethanol bath (dry ice + ethanol). We then made 10μm thick sections using a Leica cryostat, collected on Superfrost ultra plus glass slides, stored at −20°C until use. For immunohistochemistry, slides were thawn in PBS for 5 minutes and incubated in PBS, 0.3% triton X100 for 15 minutes. A step of bleaching with 1 volume 30%H2O2, 4 volumes PBS, 0.3% triton X100 (thus 6% H2O2 final concentration) was carried out for 30 minutes, after which slides were rinsed 3 times in PBS, 0.3% triton. When carrying out (see antibody list below) heat induced epitope retrieval (HIER), slides were first immersed in cold Citrate buffer HIER solution (0.1M Sodium Citrate, 0.1M Citric acid, 0.2% Tween 20, pH6.0), then incubated in Citrate buffer HIER solution preheated at 95°C, for a duration of 10 to 30 minutes (depending on primary antibody), transferred again in cold Citrate buffer HIER solution, and rinsed 3 times in PBS, 0.3% triton. The Antibody incubation steps were carried out in blocking solution containing 20% newborn calf serum, 0.3% triton X100.

Primary antibodies used are the following: Rabbit polyclonal Anti-RFP (Rockland, Cat# 600-401-379; RRID: AB_2209751); Chick IgY anti-beta-Galactosidase (Abcam, Cat# ab9361; RRID: AB_307210); Chick IgY anti-GFP (Aves, cat# GFP-1020; RRID: AB_10000240); Mouse IgG1 monoclonal anti-PAX7 (DSHB, Cat# PAX7, supernatant; RRID: AB_2299243); Mouse IgG2b monoclonal anti-MYH1 (Myosin heavy chain, type I), clone MF20 (DSHB, Cat# MF20 (bioreactor); RRID: AB_2147781); Mouse monoclonal IgG1 anti-Myogenin, clone F5D) (DSHB, via Santa-cruz biotechnology Inc. Cat# sc-12732; RRID: AB_2146602); purified Rat monoclonal anti mouse CD140a (Pdgfra), clone APA5 (BD Biosciences, Cat# 558774; RRID: AB_397117); Rat IgG1 monoclonal anti-TenascinC, clone MTn-12 (Thermofisher Scientific, Cat# MA1-26778; RRID: AB_2256026); Mouse IgG2A monoclonal anti-TCF4 (Tcf7L2), clone 6H5-3 (Merc-Millipore, Cat# 05-511; RRID: AB_309772). Antigen retrieval with Citrate buffer was done for all antibody combinations, except when using anti-GFP, anti-Pdgfra, and anti-TenascinC. We used Alexa-conjugated or Cy3/Cy5-conjugated donkey secondary antibodies against rabbit, chick and rat primary antibodies, whereas for mouse monoclonal primary antibodies, we used isotype-specific Alexa-conjugated Goat secondary antibodies (against IgG1, IgG2a; IgG2b) from Thermofisher scientific.

## Acknowledgements

We thank Angela K Zimmermann, Francesca Puppo, Marc Bartoli, Rosanna Dono, Flavio Maina, and Robert Kelly for scientific input, Carine Jiguet Jiglaire, Agnes Roure, Axel Viesel, and Nicolas Denan for generously providing some of the plasmids used to engineer the transgenic construct; Dominique Fragano for mouse genotyping; Aaron McMahon, for help with cryostat sections and immunohistochemistry; the IBDM animal house staff for mouse husbandry. Imaging was performed using the PiCSL-FBI core facility (IBDM, AMU-Marseille, France) supported by the Agence Nationale de la Recherche through the ‘Investments for the Future’ program (France-BioImaging, ANR-10-INBS-04). This work was supported by grants to FH from the AFM-Telethon (grants N°15823, N°16785 and N°20861), FSHD global research foundation (grant number Grant 14), and FSHD society (FSHS-82014-05), and by a post-doctoral FSHD society fellowship to AKZ (Grant FSHS-82012-03).

## Supplementary Information

**Figure S1:**
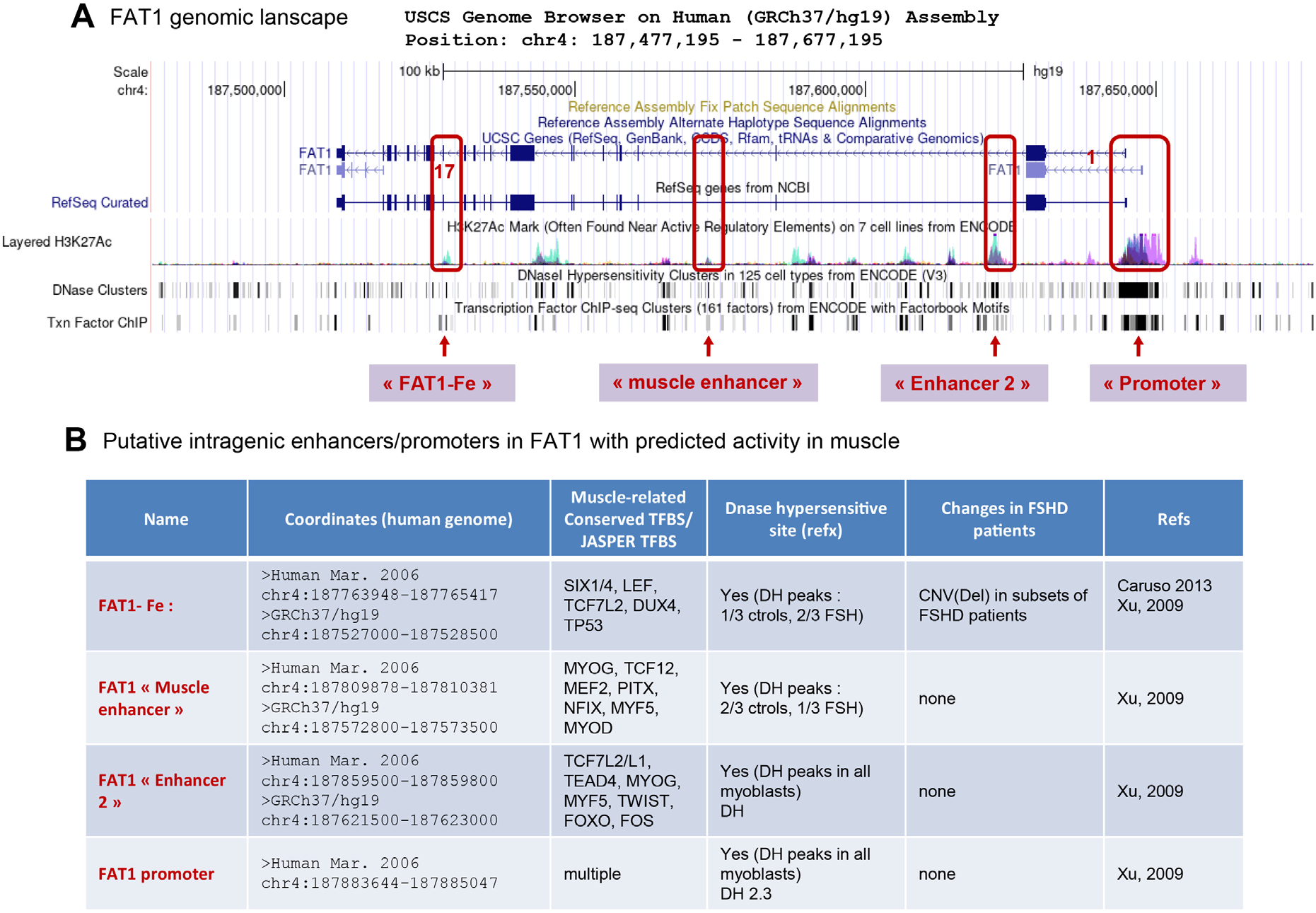
FAT1 genomic landscape: additional enhancers with predicted muscle activity. (A) Screen capture of USCS browser view of the *FAT1* genomic landscape, featuring *FAT1* Refseq with exon positions, A summary of the CHIP-Seq peaks obtained (ENCODE data) with anti-Acetylated H3K27 antibodies, indicating position of predicted enhancers, DNase hypersensitivity clusters and averaged Transcription factor CHIP-Seq peaks. The position of 4 putative cis-regulatory regions (3 intragenic FAT1 enhancers and the promoter (details in B), predicted to be active in muscle-related cell types, are highlighted, the one indicated as “FAT1-Fe” being the one deleted by FSHD-CNVs and studied here. (B) table recapitulating the names, genomic coordinates, predicted transcription factor binding sites (JASPER TFBS, whether they map with DNase hypersensitivity sites (on Ensembl browser and in human myoblasts, based on ref [68], genomic status in FSHD patients, and related references.

**Figure S2:**
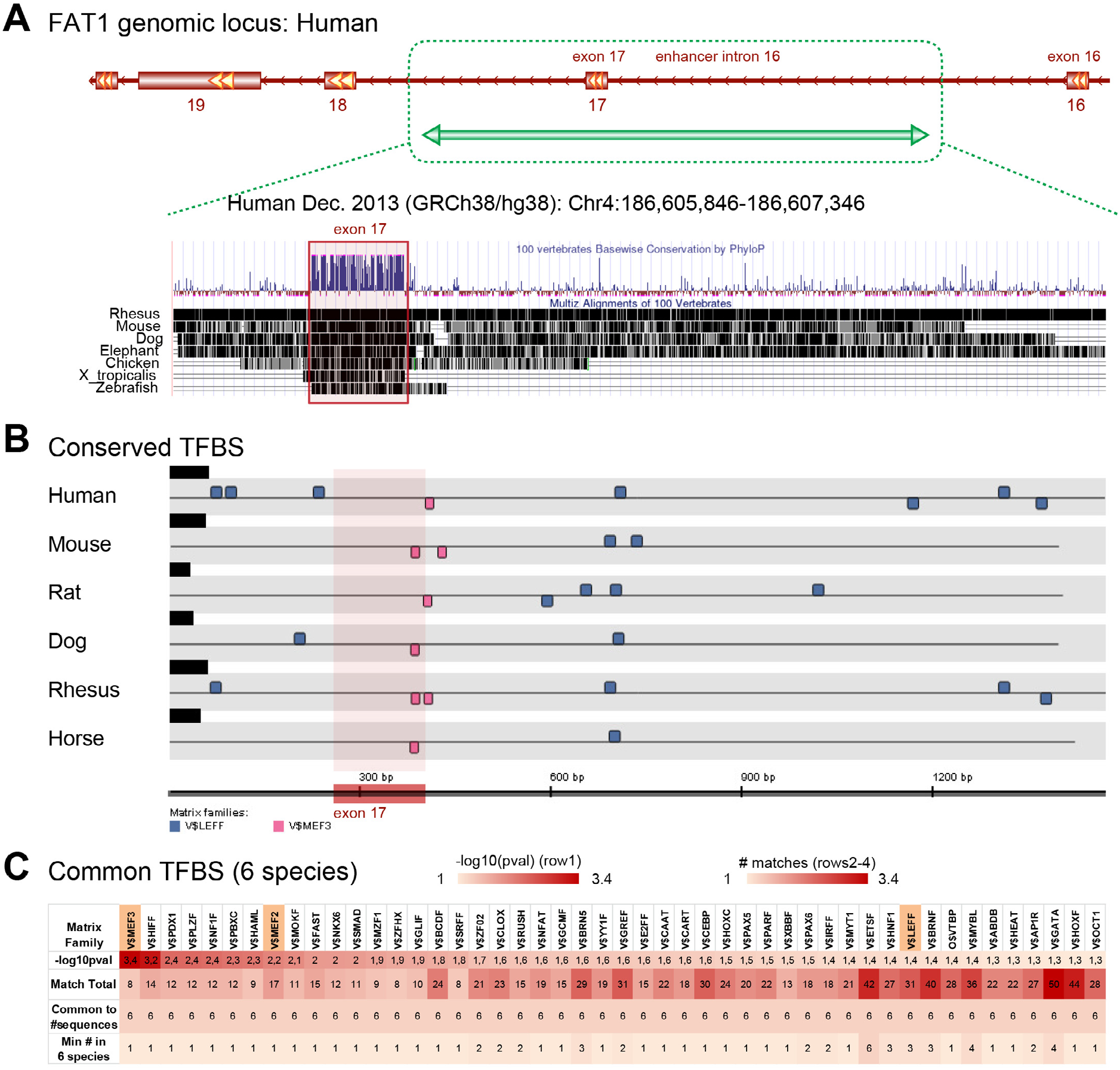
Position of previously identified copy number variants in the FAT1 genomic locus associated with FSHD. (A) Scheme of the human FAT1 locus showing exons 16 to 19, with conservation tracks from USCS, showing that the *FAT1^Fe^* sequence, deleted in the FSHD-CNV, matches a conserved region, spanning beyond the coding part corresponding to exon 17. (B) Scheme of the conservation of LEF and MEF3 (SIX1) TFBS motifs in the conserved *FAT1^Fe^* sequence across six species. (C) Heatmap of the common transcription factor binding sites (by TRANSFAC matrix family), featuring the value of (-Log10(p-value)), and to number of matches in sequences from 6 species (total, common, or minimum number).

**Figure S3:**
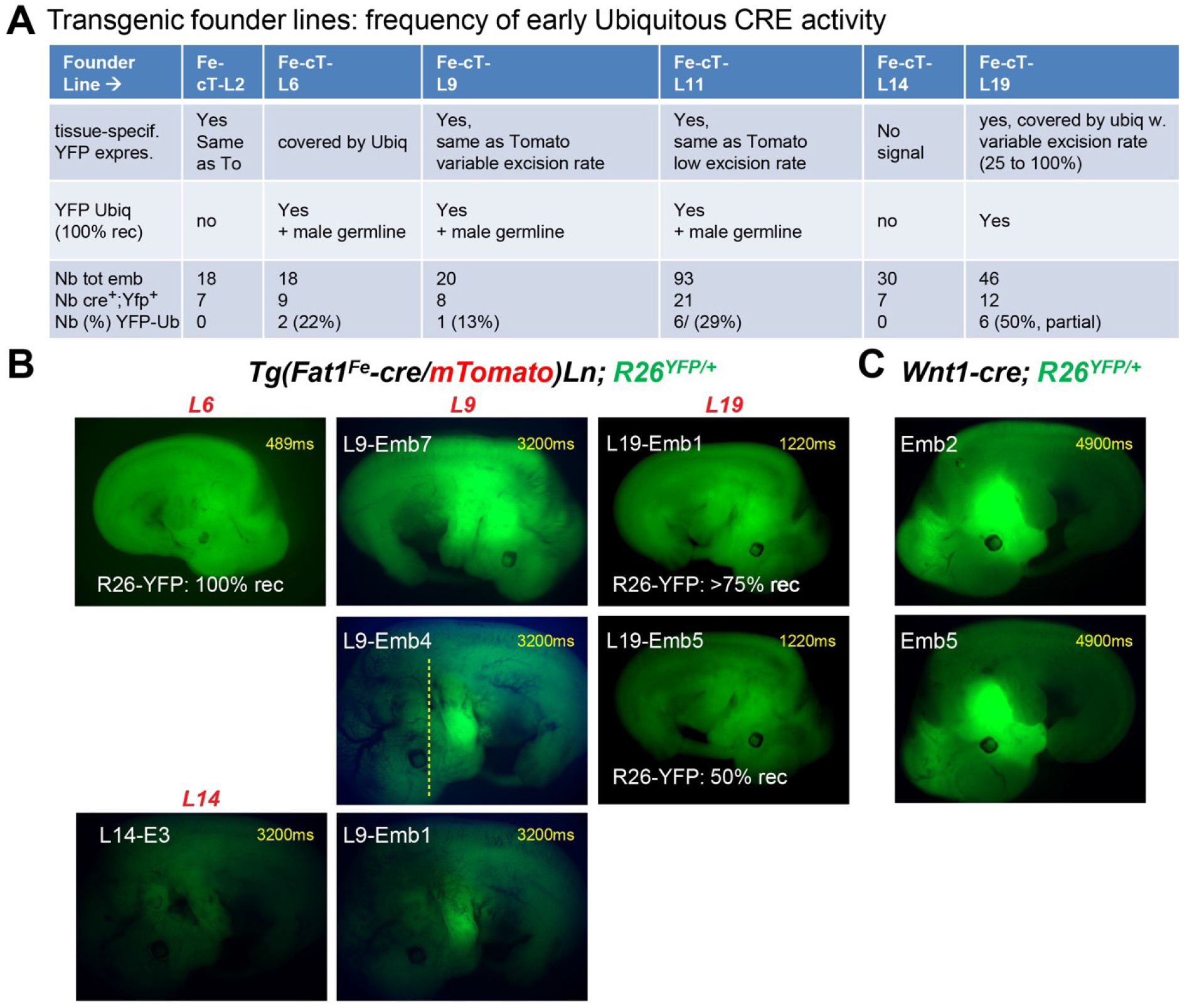
Distinction between early Ubiquitous and tissue-specific CRE activity of the founder lines. (A) Table summarizing the patterns of cre-mediated activity observed in the 6 founder lines analyzed *(Tg(Fat1Fe-cre/mTomato)Ln; R26^YFP/+^)*. As transgenic embryos carried the *R26^YFP^* reporter, fluorescence was assessed first upon embryo collection, then on embryo sections by anti-GFP immunohistochemistry. We score embryos as exhibiting tissue-specific YFP patterns when this activity matches Tomato expression (first row). We score embryos as exhibiting ubiquitous YFP (second and third row) when YFP Fluorescence is detected at high levels at collection stage (and confirmed by IHC). The last row recapitulates for each line, the total number of embryos obtained from *FF-cT-Ln* x *R26^YFP/+^* crosses, the number of *FF-cT-Ln; R26^YFP/+^* embryos (by PCR), and the number of embryos with ubiquitous YFP expression at collection (and percentage of these amongst *FF-cT-Ln; R26^YFP/+^* embryos). (B) images of direct YFP fluorescence emitted at stage of embryo collection (during fixation step) for several founder lines, with several embryos shown with variable excision rates for L9 and L19. (C) Examples of direct YFP fluorescence taken in similar conditions in 2 examples of Wnt1^cre/+^; *R26*^*YFP*/+^ embryos (with intense recombination in craniofacial neural crest-derived tissues visible without clearing).

**Figure S4:**
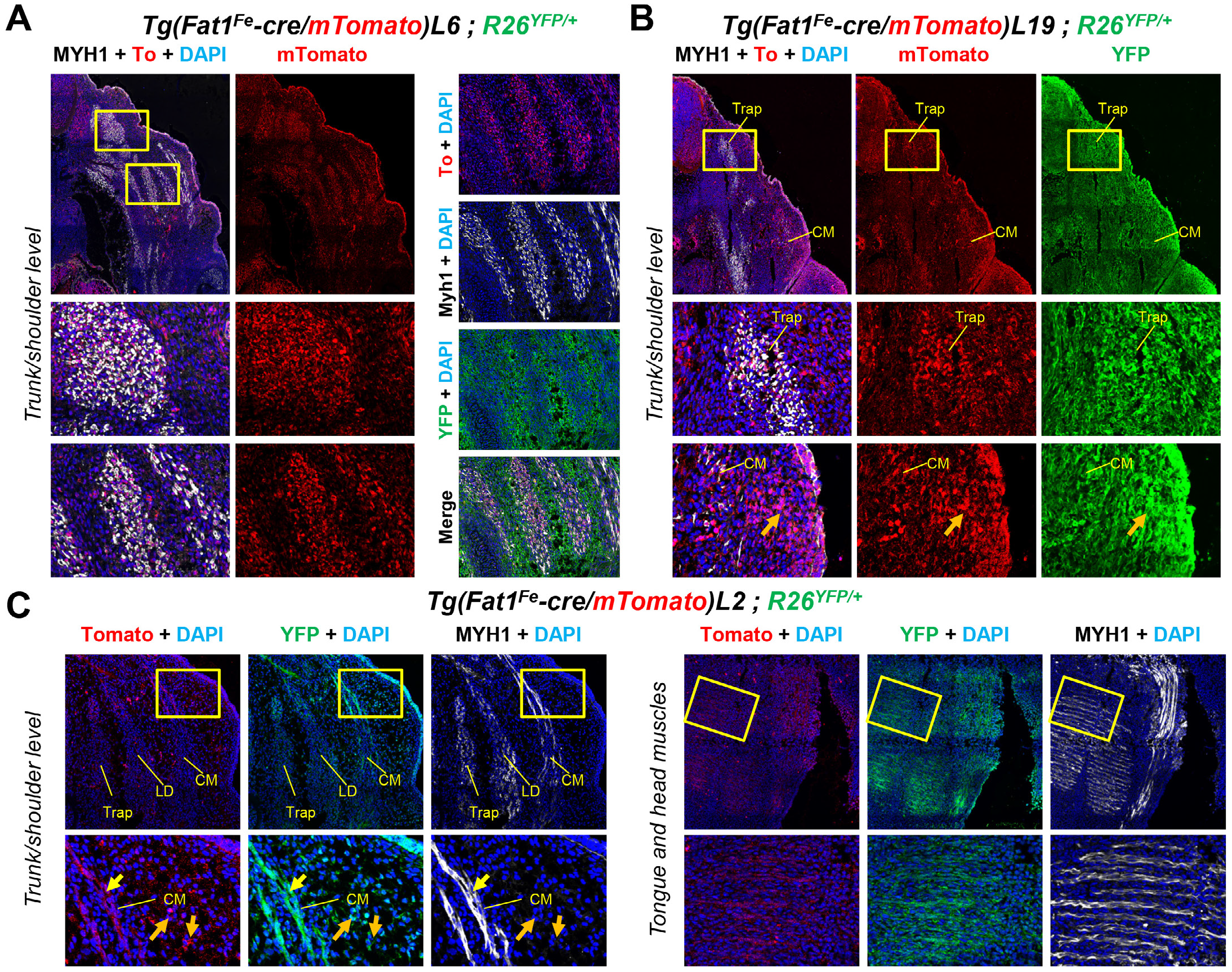
Additional examples of founder lines with muscle expression of the *FAT1^Fe^* enhancer: two of which with ubiquitous YFP expression. immunohistochemistry was performed on sections of E12.5 embryos carrying *FF-cT* lines *6* (A), *19* (B), and *2* (C) combined with *R26^YFP^*, with anti-Tomato (RFP, red), anti-MyhI (MHC, white), and anti-GFP (*R26^YFP^*, green) antibodies. The sample shown in (B) is the same as the one shown in Figure 2B, except that it is shown (horizontal flip) to illustrate ubiquitous YFP expression.

**Figure S5:**
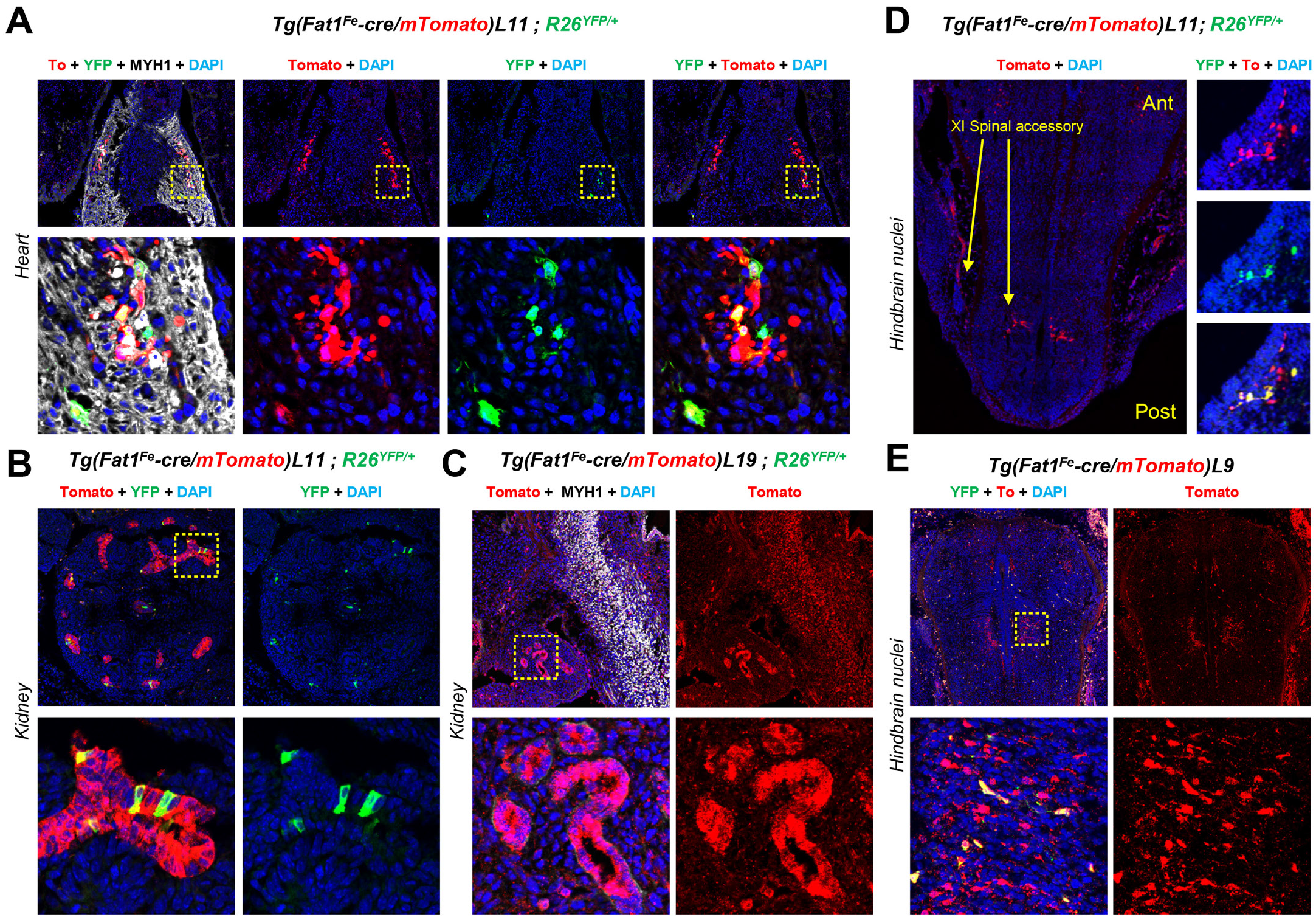
Other sites of expression/activity of the FF-cT transgene. We detected Tomato expression (A-E) and activity (via YFP expression in A, B, D, E) in the heart (A, line *FF-cT-L11)*,kidney ((B), *FF-cT-L11*, and (C), *FF-cT-L19)*, and in hindbrain nuclei ((D), *FF-cT-L11*, and (E), *FF-cT-L9*). For each line, higher panels are low magnification, and lower panels are higher magnification of the dotted square area (except for E, for which higher magnification come from a different embryo).

## References

1. Spitz F, Furlong EE: Transcription factors: from enhancer binding to developmental control. Nat Rev Genet 2012, 13(9):613–626.

2. Wittkopp PJ, Kalay G: Cis-regulatory elements: molecular mechanisms and evolutionary processes underlying divergence. Nat Rev Genet 2011, 13(1):59–69.

3. Arnoult L, Su KF, Manoel D, Minervino C, Magrina J, Gompel N, Prud’homme B: Emergence and diversification of fly pigmentation through evolution of a gene regulatory module. Science 2013, 339(6126):1423–1426.

4. D’Haene E, Bar-Yaacov R, Bariah I, Vantomme L, Van Loo S, Cobos FA, Verboom K, Eshel R, Alatawna R, Menten B et al: A neuronal enhancer network upstream of MEF2C is compromised in patients with Rett-like characteristics. Hum Mol Genet 2019, 28(5):818–827.

5. Skuplik I, Benito-Sanz S, Rosin JM, Bobick BE, Heath KE, Cobb J: Identification of a limb enhancer that is removed by pathogenic deletions downstream of the SHOX gene. Sci Rep 2018, 8(1):14292.

6. Bertorelli R, Capone L, Ambrosetti F, Garavelli L, Varriale L, Mazza V, Stanghellini I, Percesepe A, Forabosco A: The homozygous deletion of the 3’ enhancer of the SHOX gene causes Langer mesomelic dysplasia. Clin Genet 2007, 72(5):490–491.

7. Lupianez DG, Spielmann M, Mundlos S: Breaking TADs: How Alterations of Chromatin Domains Result in Disease. Trends Genet 2016, 32(4):225–237.

8. Lupianez DG, Kraft K, Heinrich V, Krawitz P, Brancati F, Klopocki E, Horn D, Kayserili H, Opitz JM, Laxova R et al: Disruptions of topological chromatin domains cause pathogenic rewiring of gene-enhancer interactions. Cell 2015, 161(5):1012–1025.

9. Zhang F, Gu W, Hurles ME, Lupski JR: Copy number variation in human health, disease, and evolution. Annu Rev Genomics Hum Genet 2009, 10:451–481.

10. Stankiewicz P, Beaudet AL: Use of array CGH in the evaluation of dysmorphology, malformations, developmental delay, and idiopathic mental retardation. Curr Opin Genet Dev 2007, 17(3):182–192.

11. Jerkovic I, Cavalli G: Understanding 3D genome organization by multidisciplinary methods. Nat Rev Mol Cell Biol 2021, 22(8):511–528.

12. Helmbacher F: Tissue-specific activities of the Fat1 cadherin cooperate to control neuromuscular morphogenesis. PLoS Biol 2018, 16(5):e2004734.

13. Caruso N, Herberth B, Bartoli M, Puppo F, Dumonceaux J, Zimmermann A, Denadai S, Lebosse M, Roche S, Geng L et al: Deregulation of the protocadherin gene FAT1 alters muscle shapes: implications for the pathogenesis of facioscapulohumeral dystrophy. PLoS Genet 2013, 9(6):e1003550.

14. Puppo F, Dionnet E, Gaillard MC, Gaildrat P, Castro C, Vovan C, Bertaux K, Bernard R, Attarian S, Goto K et al: Identification of variants in the 4q35 gene FAT1 in patients with a facioscapulohumeral dystrophy-like phenotype. Hum Mutat 2015, 36(4):443–453.

15. Mariot V, Roche S, Hourde C, Portilho D, Sacconi S, Puppo F, Duguez S, Rameau P, Caruso N, Delezoide AL et al: Correlation between low FAT1 expression and early affected muscle in facioscapulohumeral muscular dystrophy. Ann Neurol 2015, 78(3):387–400.

16. Sadeqzadeh E, de Bock CE, Thorne RF: Sleeping giants: emerging roles for the fat cadherins in health and disease. Med Res Rev 2014, 34(1):190–221.

17. Sharma P, McNeill H: Fat and Dachsous cadherins. Prog Mol Biol Transl Sci 2013, 116:215–235.

18. Saburi S, Hester I, Goodrich L, McNeill H: Functional interactions between Fat family cadherins in tissue morphogenesis and planar polarity. Development 2012, 139(10):1806–1820.

19. Gee HY, Sadowski CE, Aggarwal PK, Porath JD, Yakulov TA, Schueler M, Lovric S, Ashraf S, Braun DA, Halbritter J et al: FAT1 mutations cause a glomerulotubular nephropathy. Nat Commun 2016, 7:10822.

20. Bagherie-Lachidan M, Reginensi A, Pan Q, Zaveri HP, Scott DA, Blencowe BJ, Helmbacher F, McNeill H: Stromal Fat4 acts non-autonomously with Dchs1/2 to restrict the nephron progenitor pool. Development 2015, 142(15):2564–2573.

21. Ciani L, Patel A, Allen ND, ffrench-Constant C: Mice lacking the giant protocadherin mFAT1 exhibit renal slit junction abnormalities and a partially penetrant cyclopia and anophthalmia phenotype. Mol Cell Biol 2003, 23(10):3575–3582.

22. Helmbacher F: Astrocyte-intrinsic and -extrinsic Fat1 activities regulate astrocyte development and angiogenesis in the retina. Development 2022, 149(2).

23. Lahrouchi N, George A, Ratbi I, Schneider R, Elalaoui SC, Moosa S, Bharti S, Sharma R, Abu-Asab M, Onojafe F et al: Homozygous frameshift mutations in FAT1 cause a syndrome characterized by colobomatous-microphthalmia, ptosis, nephropathy and syndactyly. Nat Commun 2019, 10(1):1180.

24. Sugiyama Y, Shelley EJ, Badouel C, McNeill H, McAvoy JW: Atypical Cadherin Fat1 Is Required for Lens Epithelial Cell Polarity and Proliferation but Not for Fiber Differentiation. Invest Ophthalmol Vis Sci 2015, 56(6):4099–4107.

25. Badouel C, Zander MA, Liscio N, Bagherie-Lachidan M, Sopko R, Coyaud E, Raught B, Miller FD, McNeill H: Fat1 interacts with Fat4 to regulate neural tube closure, neural progenitor proliferation and apical constriction during mouse brain development. Development 2015, 142(16):2781–2791.

26. Lodge EJ, Xekouki P, Silva TS, Kochi C, Longui CA, Faucz FR, Santambrogio A, Mills JL, Pankratz N, Lane J et al: Requirement of FAT and DCHS protocadherins during hypothalamic-pituitary development. JCI Insight 2020, 5(23).

27. Peng Z, Gong Y, Liang X: Role of FAT1 in health and disease. Oncol Lett 2021, 21(5):398.

28. Frei JA, Brandenburg C, Nestor JE, Hodzic DM, Plachez C, McNeill H, Dykxhoorn DM, Nestor MW, Blatt GJ, Lin YC: Postnatal expression profiles of atypical cadherin FAT1 suggest its role in autism. Biol Open 2021, 10(6).

29. Pastushenko I, Mauri F, Song Y, de Cock F, Meeusen B, Swedlund B, Impens F, Van Haver D, Opitz M, Thery M et al: Fat1 deletion promotes hybrid EMT state, tumour stemness and metastasis. Nature 2021, 589(7842):448–455.

30. Morris LG, Kaufman AM, Gong Y, Ramaswami D, Walsh LA, Turcan S, Eng S, Kannan K, Zou Y, Peng L et al: Recurrent somatic mutation of FAT1 in multiple human cancers leads to aberrant Wnt activation. Nat Genet 2013, 45(3):253–261.

31. Cao LL, Riascos-Bernal DF, Chinnasamy P, Dunaway CM, Hou R, Pujato MA, O’Rourke BP, Miskolci V, Guo L, Hodgson L et al: Control of mitochondrial function and cell growth by the atypical cadherin Fat1. Nature 2016, 539(7630):575–578.

32. Smith TG, Van Hateren N, Tickle C, Wilson SA: The expression of Fat-1 cadherin during chick limb development. Int J Dev Biol 2007, 51(2):173–176.

33. Hou R, Liu L, Anees S, Hiroyasu S, Sibinga NE: The Fat1 cadherin integrates vascular smooth muscle cell growth and migration signals. J Cell Biol 2006, 173(3):417–429.

34. Srinivas S, Watanabe T, Lin CS, William CM, Tanabe Y, Jessell TM, Costantini F: Cre reporter strains produced by targeted insertion of EYFP and ECFP into the ROSA26 locus. BMC Dev Biol 2001, 1:4.

35. Ringel AR, Szabo Q, Chiariello AM, Chudzik K, Schöpflin R, Rothe P, Mattei AL, Zehnder T, Harnett D, Laupert V et al: Promoter repression and 3D-restructuring resolves divergent developmental gene expression in TADs (doi: 10.2139/ssrn.3947354). Cell, in press 2022.

36. Mathew SJ, Hansen JM, Merrell AJ, Murphy MM, Lawson JA, Hutcheson DA, Hansen MS, Angus-Hill M, Kardon G: Connective tissue fibroblasts and Tcf4 regulate myogenesis. Development 2011, 138(2):371–384.

37. Kardon G, Harfe BD, Tabin CJ: A Tcf4-positive mesodermal population provides a prepattern for vertebrate limb muscle patterning. Dev Cell 2003, 5(6):937–944.

38. Helmbacher F, Stricker S: Tissue cross talks governing limb muscle development and regeneration. Semin Cell Dev Biol 2020.

39. Comai G, Tajbakhsh S: Molecular and cellular regulation of skeletal myogenesis. Curr Top Dev Biol 2014, 110:1–73.

40. Lescroart F, Hamou W, Francou A, Theveniau-Ruissy M, Kelly RG, Buckingham M: Clonal analysis reveals a common origin between nonsomite-derived neck muscles and heart myocardium. Proc Natl Acad Sci U S A 2015, 112(5):1446–1451.

41. Lescroart F, Kelly RG, Le Garrec JF, Nicolas JF, Meilhac SM, Buckingham M: Clonal analysis reveals common lineage relationships between head muscles and second heart field derivatives in the mouse embryo. Development 2010, 137(19):3269–3279.

42. Nassari S, Duprez D, Fournier-Thibault C: Non-myogenic Contribution to Muscle Development and Homeostasis: The Role of Connective Tissues. Front Cell Dev Biol 2017, 5:22.

43. Vallecillo-Garcia P, Orgeur M, Vom Hofe-Schneider S, Stumm J, Kappert V, Ibrahim DM, Borno ST, Hayashi S, Relaix F, Hildebrandt K et al: Odd skipped-related 1 identifies a population of embryonic fibro-adipogenic progenitors regulating myogenesis during limb development. Nat Commun 2017, 8(1):1218.

44. Colasanto MP, Eyal S, Mohassel P, Bamshad M, Bonnemann CG, Zelzer E, Moon AM, Kardon G: Development of a subset of forelimb muscles and their attachment sites requires the ulnar-mammary syndrome gene Tbx3. Dis Model Mech 2016, 9(11):1257–1269.

45. Besse L, Sheeba CJ, Holt M, Labuhn M, Wilde S, Feneck E, Bell D, Kucharska A, Logan MPO: Individual Limb Muscle Bundles Are Formed through Progressive Steps Orchestrated by Adjacent Connective Tissue Cells during Primary Myogenesis. Cell Rep 2020, 30(10):3552-3565 e3556.

46. Collins BC, Kardon G: It takes all kinds: heterogeneity among satellite cells and fibro-adipogenic progenitors during skeletal muscle regeneration. Development 2021, 148(21).

47. Tabula Muris C: A single-cell transcriptomic atlas characterizes ageing tissues in the mouse. Nature 2020, 583(7817):590–595.

48. Rubenstein AB, Smith GR, Raue U, Begue G, Minchev K, Ruf-Zamojski F, Nair VD, Wang X, Zhou L, Zaslavsky E et al: Single-cell transcriptional profiles in human skeletal muscle. Sci Rep 2020, 10(1):229.

49. Oprescu SN, Yue F, Qiu J, Brito LF, Kuang S: Temporal Dynamics and Heterogeneity of Cell Populations during Skeletal Muscle Regeneration. iScience 2020, 23(4):100993.

50. Giordani L, He GJ, Negroni E, Sakai H, Law JYC, Siu MM, Wan R, Corneau A, Tajbakhsh S, Cheung TH et al: High-Dimensional Single-Cell Cartography Reveals Novel Skeletal Muscle-Resident Cell Populations. Molecular Cell 2019, 25 March 2019.

51. Arostegui M, Wilder Scott R, Bose K, Michael Underhill T: Cellular taxonomy of Hic1(+) mesenchymal progenitor derivatives in the limb: from embryo to adult. Nat Commun 2022, 13(1):4989.

52. Schatzl T, Kaiser L, Deigner HP: Facioscapulohumeral muscular dystrophy: genetics, gene activation and downstream signalling with regard to recent therapeutic approaches: an update. Orphanet J Rare Dis 2021, 16(1):129.

53. DeSimone AM, Pakula A, Lek A, Emerson CP, Jr.: Facioscapulohumeral Muscular Dystrophy. Compr Physiol 2017, 7(4):1229–1279.

54. Lemmers RJ, van der Vliet PJ, Klooster R, Sacconi S, Camano P, Dauwerse JG, Snider L, Straasheijm KR, van Ommen GJ, Padberg GW et al: A unifying genetic model for facioscapulohumeral muscular dystrophy. Science 2010, 329(5999):1650–1653.

55. Hendrickson PG, Dorais JA, Grow EJ, Whiddon JL, Lim JW, Wike CL, Weaver BD, Pflueger C, Emery BR, Wilcox AL et al: Conserved roles of mouse DUX and human DUX4 in activating cleavage-stage genes and MERVL/HERVL retrotransposons. Nat Genet 2017, 49(6):925–934.

56. De Iaco A, Planet E, Coluccio A, Verp S, Duc J, Trono D: DUX-family transcription factors regulate zygotic genome activation in placental mammals. Nat Genet 2017, 49(6):941–945.

57. Hewitt JE: Loss of epigenetic silencing of the DUX4 transcription factor gene in facioscapulohumeral muscular dystrophy. Hum Mol Genet 2015, 24(R1):R17–23.

58. Gaillard MC, Broucqsault N, Morere J, Laberthonniere C, Dion C, Badja C, Roche S, Nguyen K, Magdinier F, Robin JD: Analysis of the 4q35 chromatin organization reveals distinct long-range interactions in patients affected with Facio-Scapulo-Humeral Dystrophy. Sci Rep 2019, 9(1):10327.

59. van den Boogaard ML, Lemmers RJ, Balog J, Wohlgemuth M, Auranen M, Mitsuhashi S, van der Vliet PJ, Straasheijm KR, van den Akker RF, Kriek M et al: Mutations in DNMT3B Modify Epigenetic Repression of the D4Z4 Repeat and the Penetrance of Facioscapulohumeral Dystrophy. Am J Hum Genet 2016, 98(5):1020–1029.

60. Lemmers RJ, Tawil R, Petek LM, Balog J, Block GJ, Santen GW, Amell AM, van der Vliet PJ, Almomani R, Straasheijm KR et al: Digenic inheritance of an SMCHD1 mutation and an FSHD-permissive D4Z4 allele causes facioscapulohumeral muscular dystrophy type 2. Nat Genet 2012, 44(12):1370–1374.

61. Scionti I, Greco F, Ricci G, Govi M, Arashiro P, Vercelli L, Berardinelli A, Angelini C, Antonini G, Cao M et al: Large-scale population analysis challenges the current criteria for the molecular diagnosis of fascioscapulohumeral muscular dystrophy. Am J Hum Genet 2012, 90(4):628–635.

62. Scionti I, Fabbri G, Fiorillo C, Ricci G, Greco F, D’Amico R, Termanini A, Vercelli L, Tomelleri G, Cao M et al: Facioscapulohumeral muscular dystrophy: new insights from compound heterozygotes and implication for prenatal genetic counselling. J Med Genet 2012, 49(3):171–178.

63. Jones TI, Chen JC, Rahimov F, Homma S, Arashiro P, Beermann ML, King OD, Miller JB, Kunkel LM, Emerson CP, Jr. et al: Facioscapulohumeral muscular dystrophy family studies of DUX4 expression: evidence for disease modifiers and a quantitative model of pathogenesis. Hum Mol Genet 2012, 21(20):4419–4430.

64. de Greef JC, Krom YD, den Hamer B, Snider L, Hiramuki Y, van den Akker RFP, Breslin K, Pakusch M, Salvatori DCF, Slutter B et al: Smchd1 haploinsufficiency exacerbates the phenotype of a transgenic FSHD1 mouse model. Hum Mol Genet 2018, 27(4):716–731.

65. Sacconi S, Lemmers RJ, Balog J, van der Vliet PJ, Lahaut P, van Nieuwenhuizen MP, Straasheijm KR, Debipersad RD, Vos-Versteeg M, Salviati L et al: The FSHD2 gene SMCHD1 is a modifier of disease severity in families affected by FSHD1. Am J Hum Genet 2013, 93(4):744–751.

66. Park HJ, Lee W, Kim SH, Lee JH, Shin HY, Kim SM, Park KD, Choi YC: FAT1 Gene Alteration in Facioscapulohumeral Muscular Dystrophy Type 1. Yonsei Med J 2018, 59(2):337–340.

67. Gaildrat P, Killian A, Martins A, Tournier I, Frebourg T, Tosi M: Use of splicing reporter minigene assay to evaluate the effect on splicing of unclassified genetic variants. Methods Mol Biol 2010, 653:249–257.

68. Xu X, Tsumagari K, Sowden J, Tawil R, Boyle AP, Song L, Furey TS, Crawford GE, Ehrlich M: DNaseI hypersensitivity at gene-poor, FSH dystrophy-linked 4q35.2. Nucleic Acids Res 2009, 37(22):7381–7393.

69. Kvon EZ, Waymack R, Gad M, Wunderlich Z: Enhancer redundancy in development and disease. Nat Rev Genet 2021, 22(5):324–336.

70. Osterwalder M, Barozzi I, Tissieres V, Fukuda-Yuzawa Y, Mannion BJ, Afzal SY, Lee EA, Zhu Y, Plajzer-Frick I, Pickle CS et al: Enhancer redundancy provides phenotypic robustness in mammalian development. Nature 2018, 554(7691):239–243.

71. Roure A, Rothbacher U, Robin F, Kalmar E, Ferone G, Lamy C, Missero C, Mueller F, Lemaire P: A multicassette Gateway vector set for high throughput and comparative analyses in ciona and vertebrate embryos. PLoS One 2007, 2(9):e916.

72. Pennacchio LA, Ahituv N, Moses AM, Prabhakar S, Nobrega MA, Shoukry M, Minovitsky S, Dubchak I, Holt A, Lewis KD et al: In vivo enhancer analysis of human conserved non-coding sequences. Nature 2006, 444(7118):499–502.

73. Leighton PA, Mitchell KJ, Goodrich LV, Lu X, Pinson K, Scherz P, Skarnes WC, Tessier-Lavigne M: Defining brain wiring patterns and mechanisms through gene trapping in mice. Nature 2001, 410(6825):174–179.

74. Danielian PS, Muccino D, Rowitch DH, Michael SK, McMahon AP: Modification of gene activity in mouse embryos in utero by a tamoxifen-inducible form of Cre recombinase. Curr Biol 1998, 8(24):1323–1326.

